# The dietary fiber guar gum ameliorates experimental autoimmune encephalomyelitis via attenuated Th1 activation and differentiation

**DOI:** 10.1101/2022.08.04.502686

**Authors:** Naomi M Fettig, Hannah G Robinson, Jessica R Allanach, Katherine M Davis, Rachel L Simister, Elsie J Wang, Ava J Sharon, Ye Jiayu, Sarah J Popple, Jung Hee Seo, Deanna L Gibson, Sean A Crowe, Marc S Horwitz, Lisa C Osborne

**Author notes:** Lead Contact: Lisa Osborne.

## Abstract

Dietary fibers are potent modulators of immune responses that can restrain inflammation in multiple disease contexts. However, dietary fibers encompass a biochemically diverse family of carbohydrates, and it remains unknown how different fiber sources influence immunity. In a head-to-head comparison of four different high-fiber diets, we demonstrate a unique and potent ability of guar gum to reduce neuroinflammation in experimental autoimmune encephalomyelitis (EAE), a T cell-mediated mouse model of multiple sclerosis. CD4^+^ T cells from guar gum-fed mice have blunted Th1-skewing, reduced migratory capacity, and limited activation and proliferative capabilities. These changes are not explained by guar gum-specific alterations to the microbiota at the 16S rRNA level, nor by specific alterations in short chain fatty acids. These findings demonstrate specificity in the host response to fiber sources, and define a new pathway of fiber-induced CD4^+^ T cell immunomodulation that protects against pathologic neuroinflammation.

## Introduction

The increasing incidence of autoimmune, inflammatory, and metabolic disorders across resource-rich countries in the last few decades is indicative of environmental, rather than genetic, changes within affected populations. Dietary choices are one environmental factor that have been suggested to contribute, and mechanistic studies in pre-clinical disease models support the assertion that Westernized diets (high fat, salt, iron, sugar) promote inflammatory immune profiles. In contrast, traditional/cultural diets may provide protective effects (Christ et al., 2019; O’Keefe et al., 2015; Sonnenburg and Sonnenburg, 2019). The intestinal microbiota is exquisitely sensitive to dietary intake, and dietary changes can alter microbial community composition and function, including microbe-derived metabolites that impact host physiology. Indeed, gut dysbiosis is consistently linked with various inflammatory and autoimmune diseases (Belkaid and Hand, 2014; de Oliveira et al., 2017; Xu et al., 2019). The relationship between diet, microbiota-derived metabolites, and the immune system implicates dietary substrates as potent sources of immunomodulation that could favor development of inflammatory diseases or be exploited for therapeutic purposes.

Multiple sclerosis (MS) is an autoimmune disease of the central nervous system that results in neuronal demyelination and subsequent neurodegeneration that manifests in cognitive and/or motor dysfunction in affected individuals. The immunopathology of MS is complex, including recruitment of inflammatory macrophages, autoreactive B cells, CD8^+^ T cells, and CD4^+^ T helper (Th) 1 and Th17 T cells to the CNS (Dendrou et al., 2015). Simultaneously, CD4^+^ regulatory T cells (Tregs) and other immunomodulatory populations are recruited to limit inflammation (Ruiz et al., 2019). Like many chronic inflammatory disorders, MS risk is best explained by a multifactorial model that accounts for genetic susceptibility plus environmental exposures, including geographic location, nutrition, infection history, and smoking (Baranzini and Oksenberg, 2017; Olsson et al., 2016). Many of these environmental factors impact the microbiota, and studies with independent cohorts have demonstrated that the microbiota of people with MS differs from healthy controls (Chen et al., 2016; Jangi et al., 2016; Miyake et al., 2015). These microbial communities appear to contribute to MS pathophysiology, as transfer of MS microbiomes into mice enhances MS-like disease (Berer et al., 2017; Cekanaviciute et al., 2017).

Diet plans that include increased dietary fiber intake are common among people with MS (Beckett et al., 2019), and several studies have shown benefits of high fiber intake to quality of life and well-being in MS patients (Evers et al., 2021; Fitzgerald et al., 2018). Although there are limited numbers of clinical trials evaluating dietary intervention in people with MS, an intervention of increased vegetable intake and limited protein intake (in essence a high-fiber diet) resulted in reduced disability scores, fewer relapses, and reduced circulating CD4^+^IL-17A^+^ T cells compared to participants that remained on a Westernized diet (Saresella et al., 2017).

Mechanistic understanding of how dietary alterations contribute to reduced inflammation and dampened autoimmunity, however, remains limited to animal models. Dietary fiber has anti-inflammatory effects in various infection models, including influenza (Trompette et al., 2018), respiratory syncytial virus (Antunes et al., 2019), *Clostridium difficile* (Hryckowian et al., 2018), and in models of emphysema (Jang et al., 2021), rheumatoid arthritis (Bai et al., 2021), systemic lupus erythematosus (Zegarra-Ruiz et al., 2019), and type 1 diabetes (Zou et al., 2021). In the context of EAE, the effects of the soluble fibers pectin and inulin conferred only modest protection, even in the presence of increased short-chain fatty acid (SCFA) production and numbers of Tregs (Mizuno et al., 2017; Park et al., 2019). To date, supplementation with the non-fermentable dietary fiber cellulose has provided the most promising data as a protective intervention in EAE (Berer et al., 2018). These data suggest that fermentable vs non-fermentable fiber sources differentially affect the host, yet it remains unknown whether fermentable dietary fibers are interchangeable in microbiota-immune system cross-talk. To address this, we performed a head-to-head comparison of four different fermentable fiber sources to assess whether they exert similar or differential effects in the context of EAE.

Here, we demonstrate that dietary supplementation with distinct single sources of soluble fermentable fiber (inulin, pectin, resistant starch, or guar gum) differentially impact CD4^+^ T cell function and EAE outcome. The fiber guar gum was uniquely protective, and this was associated with cell intrinsic defects in CD4^+^ Th1 activation and impaired CD4^+^ T cell migratory potential, resulting in reduced neuroinflammation following EAE immunization. Interestingly, Th17 differentiation appeared unaffected. Dietary fiber supplementation drove changes in microbiota community composition, but we did not detect a guar gum-specific microbial signature nor unique alterations in SCFA levels. Collectively, these data demonstrate previously unappreciated specificity in host-microbiota pathways of fermentable fiber utilization, with distinct immunomodulatory effects that could be leveraged for targeted support of inflammatory disorders.

## Results

### Dietary supplementation with guar gum, but not other soluble fibers, ameliorates EAE onset, severity, and CNS infiltration

To test the hypothesis that autoimmune neuroinflammation is responsive to dietary fiber intake, we exposed cohorts of mice to standard 5% cellulose (control) fiber diet, diet entirely lacking dietary fiber (0% fiber), or diets enriched (30%) with a single fiber source of resistant starch, inulin, pectin, or guar gum (macronutrient information in Table S1) for two weeks prior to EAE induction (Figure 1A). In contrast to findings in other inflammatory disease contexts that showed increased inflammation with fiber-deficient diets (Desai et al., 2016; Neumann et al., 2021; Shen et al., 2021; Tan et al., 2016; Trompette et al., 2014), but consistent with previous findings in EAE (Mizuno et al., 2017), a zero-fiber diet did not significantly affect EAE onset, incidence, or severity (Figure 1B-D).

**Figure 1.**
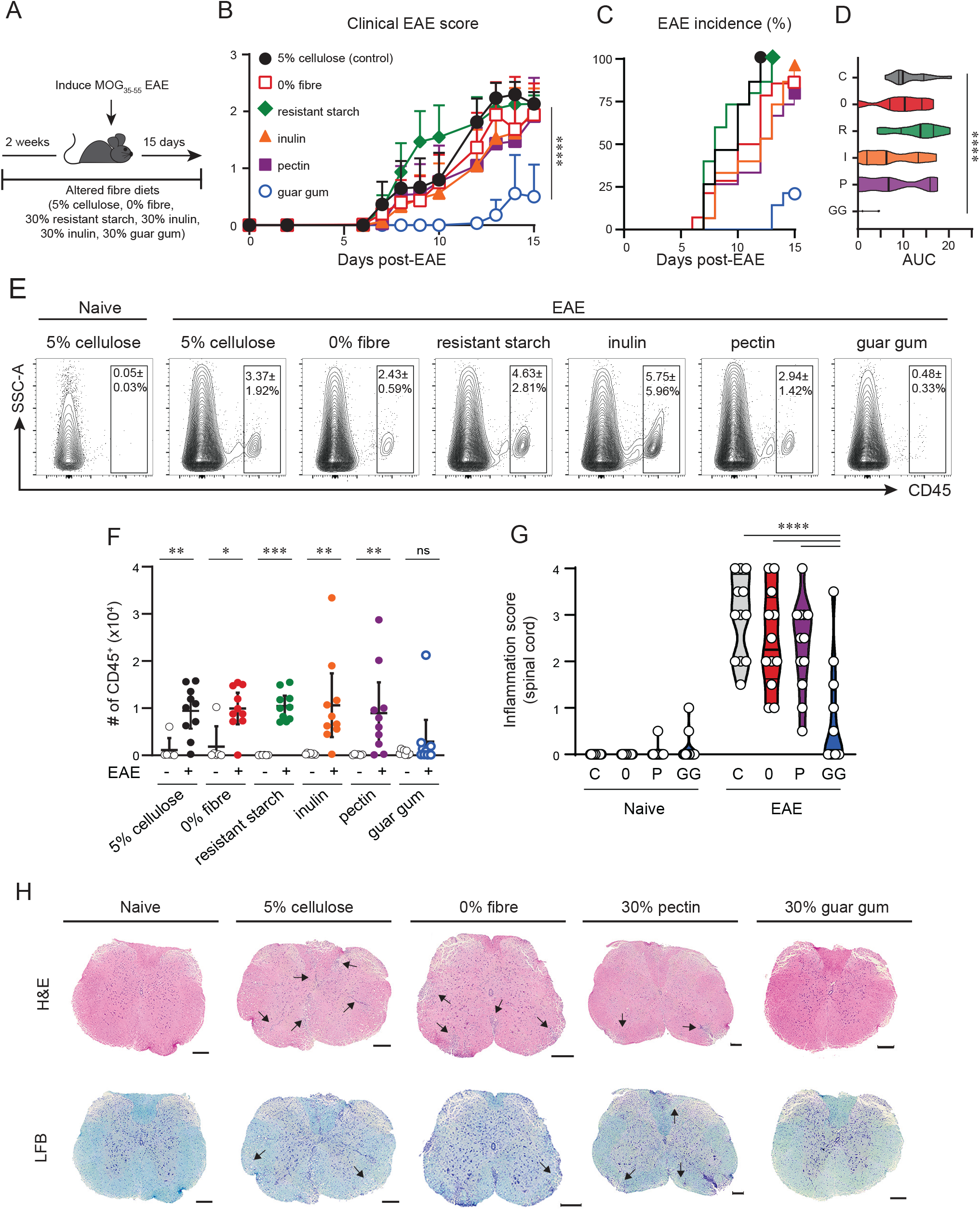
Guar gum supplementation ameliorates EAE onset, severity, and spinal cord infiltration. **A)** Mice are fed altered-fiber diets (5% cellulose (control, C), 0% fiber (0), 30% resistant starch (R), inulin (I), pectin (P), or guar gum (GG)) from 2 weeks prior to MOG_35-55_ immunization until experimental endpoint on day (d)15 post-EAE. **B-D)** Clinical EAE scores, incidence, and area under the curve (AUC) of mice on each diet (n=13-15 per group). Results pooled from 3 independent experiments. **E)** Concatenated flow plots from one representative experiment of spinal cord (SC)-infiltrating CD45^hi^ cells. Proportion of live cells shown as mean ± SD. **F)** Number of infiltrating CD45^hi^ cells in the SC of naïve and EAE-induced mice. **G)** Quantification of cellular infiltration in SC sections. **H)** Representative serial sections of SC histology by hematoxylin and eosin (H&E) and luxol fast blue (LFB). Naïve sample from 5% cellulose-fed mouse without EAE. Arrows depict regions of cellular infiltration or demyelination. Scale bar = 200 μm. Data shown as mean ± 95% CI (B,F) or violin plots depicting median and 1^st^ and 3^rd^ quartiles (D,G). Stats by 2-way ANOVA with Tukey’s multiple comparisons test (A; guar gum vs. each diet individually d15 post-EAE,G), Kruskal-Wallis with Dunn’s multiple comparisons test (D;5% cellulose vs. each other diet), or multiple Mann-Whitney tests with Holm-Šídák multiple comparisons test (F). See also Figure S1.

The fermentable fibers resistant starch, inulin, pectin, and guar gum differ in their molecular structures, but little is known about how these differences ultimately impact immune function. Over a 15-day monitoring period, dietary supplementation with resistant starch, inulin, or pectin offered no significant protection in terms of EAE onset, severity, or incidence. However, compared to control- and other fiber-enriched diets, guar gum supplementation significantly delayed symptom onset, limited disease incidence, and reduced cumulative disease severity (Figure 1B-D). The protective effect of dietary guar gum was long-lasting: at day (d) 25 post-EAE induction, 40% of guar gum-fed mice remained EAE-free, although the maximal clinical severity was not impeded in guar gum-fed mice that did develop disease (Figure S1A-C).

Flow cytometric assessment of CNS inflammation at d15 post-EAE induction revealed a significant EAE-induced enrichment of CD45^+^ cells into the brain and spinal cord (SC) of mice fed control, zero-fiber, resistant starch-, inulin-, or pectin-enriched diets, that was not apparent in guar gum-fed mice (Figure 1E-F, Figure S1D-E). Supporting these findings, blinded scoring of histological sections revealed reduced inflammation in the brain and SC of guar gum fed mice, with minimal SC demyelination compared to control and pectin-enriched diets (Figure 1G-H, Figure S1F-G). Collectively, these data indicate that dietary fiber content can differentially influence inflammatory disease outcome and highlight a uniquely protective effect of guar gum that is associated with reduced recruitment of immune cells into the CNS following EAE induction.

### Generalized reductions in intestinal microbial diversity and increased SCFA in response to high-fiber diets

The microbiota is exquisitely sensitive to diet and is an important modulator of immune function. We performed 16S rRNA sequencing to profile fecal microbial communities of mice fed control diet, diets lacking fiber, and diets supplemented with resistant starch, inulin, pectin, or guar gum. The fecal microbial communities of mice that had been fed experimental diets for two weeks clustered by diet and differed significantly from each other (Figure 2A, Figure S2A, Table S2). Notably, deprivation or supplementation with a single source of dietary fiber were uniformly associated with reduced microbial a-diversity (Figure 2B), suggesting that a diet with a specific complex carbohydrate source can generate a ‘nutritional niche’ that selects for a limited number of microbes that can utilize the available carbohydrate structures.

**Figure 2.**
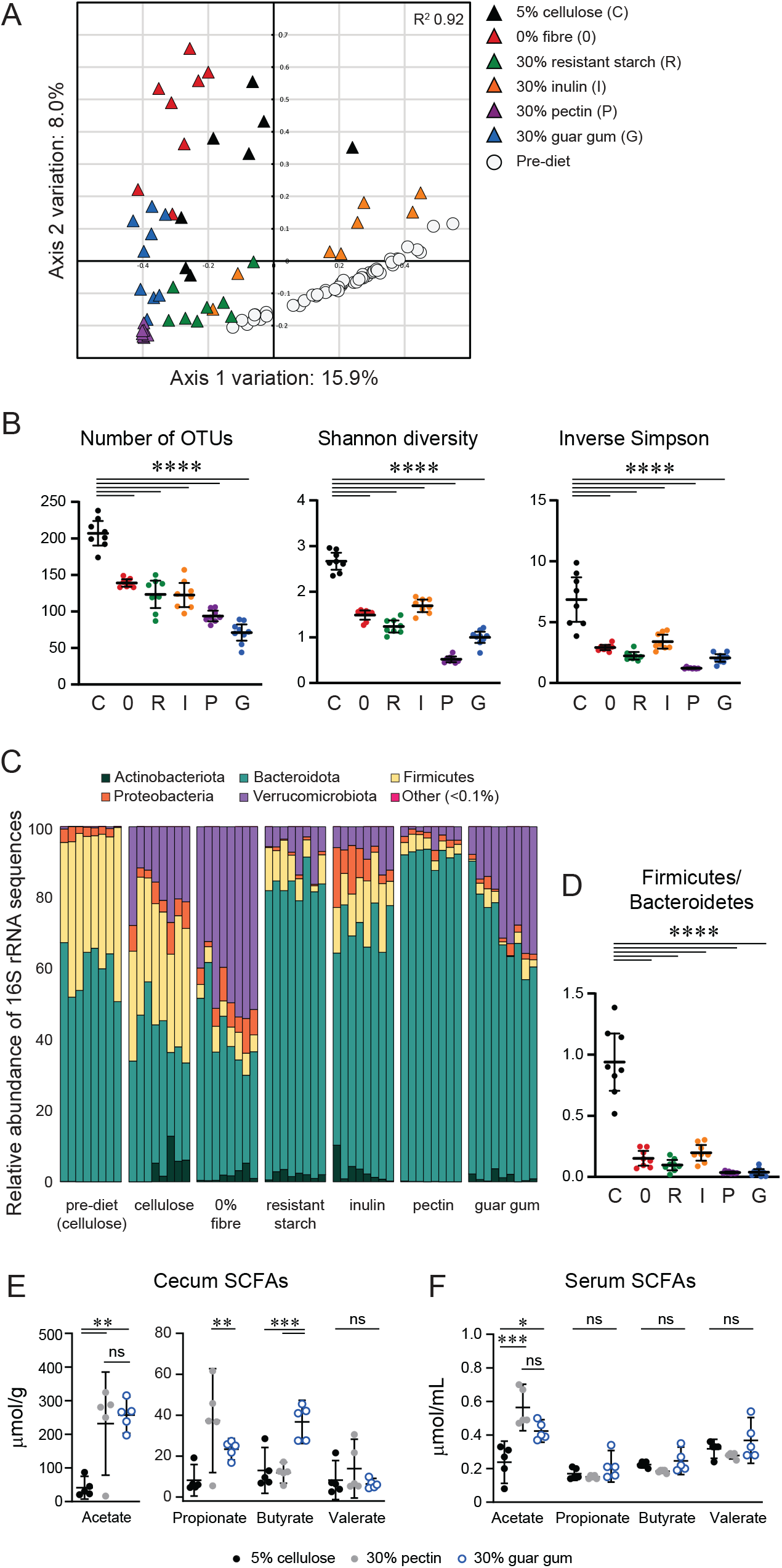
Dietary fiber supplementation reduces intestinal microbial diversity. **A)** PCoA of 16S rRNA sequences of fecal microbiota from mice fed 5% cellulose (C), 0% fiber (0), 30% resistant starch (R), 30% inulin (I), 30% pectin (P), and 30% guar gum (G) for 2 weeks. Pre-diet samples from mice upon arrival in facility prior to randomization between cages. **B)** α-diversity of fecal 16S sequences. **C)** Phylum-level abundance of 16S rRNA sequences shown for individual mice. Representatives from pre-diet group shown. **D)** Ratio of Firmicutes to Bacteroidetes. **E,F)** Short-chain fatty acid (SCFA) quantification in the cecum and serum. Data shown as mean ± 95% CI. Stats by one-way ANOVA with Tukey’s multiple comparisons test (B,D-F). See also Figure S2 and Table S1-S3.

Phylum-level 16S rRNA sequences revealed high-level changes to the microbial community structure of both zero- and high-fiber diets. In accordance with previous findings (Desai et al., 2016), differences between the 5% control diet and zero-fiber diet were driven by an increased relative abundance of Verrucomicrobia (Figure 2C), which was supported by identification of OTU_0003 (*Akkermansia* sp.) as an indicator species for the zero-fiber diet by linear discriminatory analysis effect size (LEfSe) (Table S3A). Interestingly, all high-fiber diets were associated with significantly reduced Firmicutes: Bacterioidetes ratio compared to controls (Figure 2D), which is consistent with previous studies of inulin or pectin supplementation and can be visualized in the β-diversity representation determined by the EnvFit function (Figure S2B) (Jang et al., 2021; Trompette et al., 2014, 2018). LEfSe analysis of fecal microbial communities from guar gum-fed mice compared to all other diets (All Diets, containing samples from control, zero-fiber, resistant starch-, inulin- and pectin-fed-mice) identified three OTUs (OTU_0003, *Akkermansia*; OTU0071, *Lachnospiraceae_UCG-001*; and OTU0231, a low-abundance (<0.05%) *Akkermansia*) as being over-represented in guar gum-fed mice compared to the All Diet group (Figure S2C-D, Table S2B). However, when the diets were disaggregated, the signatures of selective over- or under-representation in taxa identified in the All Diet comparison were lost apart from OTU_0231 (Figure S2E, Table S3A). Indicator value analysis (IndVal) (Dufrêne and Legendre, 1997), an alternative method to identify taxa specifically indicative of a treatment group, identified 4 OTUs of the Lachnospiraceae family (Table S3C) associated with guar gum-fed mice compared to all other diets. However, similar to LEfSe analysis, these alterations in low-abundance taxa appear to be driven by variation in individual mice rather than within the entire group (Table S3C). Collectively, these data indicate that the overall shift in microbial community composition conferred by guar gum supplementation may be more meaningful than the emergence (or loss) of individual taxa.

To address one of the functional outputs of microbial community function commonly associated with fiber-enriched dietary interventions, we measured the abundance of the SCFAs acetate, propionate, butyrate, and valerate in the cecal contents and serum of mice fed control, pectin-rich, or guar gum-rich diets. We chose pectin as our comparator since it has previously been shown to reduce EAE severity (Mizuno et al., 2017) and because the two diets could be evenly matched for macronutrient content (Table S1). Consistent with earlier studies, mice receiving a pectin-supplemented diet had elevated concentrations of cecal SCFAs, including acetate and propionate, compared to control (Figure 2E) (Lewis et al., 2019; Mizuno et al., 2017; Trompette et al., 2014). In guar gum-fed mice, cecal acetate and butyrate were also elevated (Figure 2E), and total cecal SCFAs were similar between guar gum- and pectin-fed mice (Figure S2F). The increase in butyrate in guar gum-fed mice may be related to elevated relative abundance of Lachnospiraceae OTUs detected by IndVal (Table S3C), members of which are known butyrate producers, although this function varies widely within the family (Vacca et al., 2020). Notably, the only SCFA elevated in the serum was acetate, and it was present at similar levels in pectin- and guar gum-fed mice (Figure 2F). Thus, although a guar gum-enriched diet stimulated production of SCFAs that have previously been shown to be immunoregulatory and protective in EAE (Luu et al., 2019; Mizuno et al., 2017; Park et al., 2019), the elevated cecal and circulating concentrations cannot account for the dramatic differences in clinical outcome between pectin and guar gum.

### Reduced infiltration of IFNγ-producing CD4^+^ T cells into the CNS following guar gum supplementation

Histologic and flow cytometric findings indicate reduced neuroinflammation in guar gum-fed mice compared to controls or other high fiber diets (Figure 1E-H). We used flow cytometry to determine which cell types were recruited or excluded from the CNS. At d15 post-EAE, there was no difference in the number of recruited B220^+^ B cells, but a marked reduction in the number of T cells recovered from the SC of guar gum-fed mice compared to all other diets (Figure S3A, B), suggesting that T cells may be preferentially excluded from the CNS of guar gum-fed mice.

MOG_35-55_/CFA-induced EAE in C57BL/6 mice is generally considered a CD4^+^ T cell-mediated autoimmune disease driven by IFNγ-producing Th1 and IL-17A-producing Th17 cells that can be restrained by recruitment of Foxp3^+^ Tregs (Simmons et al., 2013). Further assessment of the CNS-infiltrating CD4^+^ T cell populations demonstrated a stark reduction in the frequency and number of IFNγ-producing cells in the CNS of guar gum-fed mice (Figure 3A-C, Figure S3C-D). In the few SC-infiltrating Th1 cells isolated from guar gum-fed mice, there was a trend toward reduced IFNγ expression compared to other diets (measured by median fluorescence intensity (MFI)) (Figure 3D, S3E). In contrast, there was no change in the frequency, or MFI, of IL-17A^+^ cells (Figure 3A-D, Figure S3C-E), or in the frequency or number of dual-producing IFNγ^+^ IL-17A^+^ cells (Figure S3C-D, Figure S3F-G). Finally, the frequency and number of Foxp3^+^ CD4^+^ Tregs were similar in the SC across all diets (Figure 3E-G, Figure S3H-I) and there was no difference in the ratio of inflammatory effector CD4^+^ T cells to regulatory T cells (T_eff_/T_reg_) across all diets (Figure 3H, Figure S3J), suggesting that the selective protection from EAE seen in guar gum-fed mice is not driven by an influx of immunosuppressive Tregs. Collectively, these data suggest a specific impairment in the accumulation of pathogenic Th1 cells in the CNS following guar gum supplementation.

**Figure 3.**
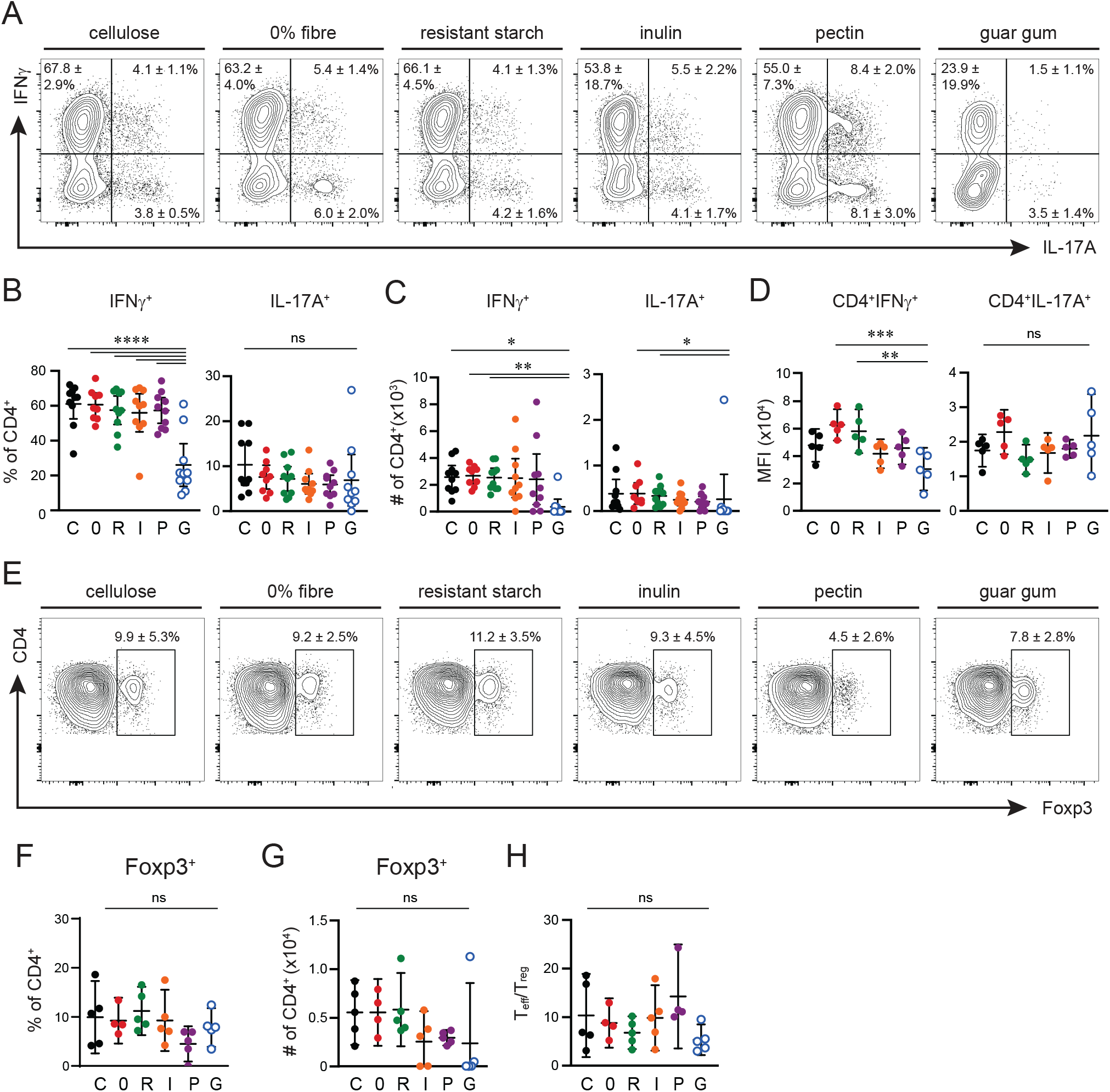
Reduced infiltration of Th1s into the spinal cord following guar gum supplementation. **A)** Concatenated flow plot from one representative experiment following PMA/ionomycin stimulation in spinal cord (SC) cells isolated from mice fed 5% cellulose (C), 0% fiber, (0), resistant starch (R), inulin (I), pectin (P), or guar gum (G) d15 post-EAE. IFNγ and IL-17A expression of live CD45^+^TCRβ^+^CD4^+^ cells shown as mean ± SD. **B,C)** Proportion and number of IFNγ^+^ or IL-17A^+^ CD4 T cells in SC. **D)** Median fluorescent intensity (MFI) of IFNγ and IL-17A of CD4^+^ T cells. **E)** Concatenated flow plot showing representative Foxp3 expression of live CD45^+^TCRß^+^CD4^+^ cells isolated from the SC at d15 post-EAE, with mean ± SD of each group. **F,G)** Proportion and number of Foxp3^+^ CD4 T cells in SC. **H)** Ratio of cytokine-expressing (IFNγ^+^, IL-17A^+^, IFNγ^+^IL-17A^+^) effector CD4^+^ T cells to Foxp3-expressing CD4^+^ Treg in SC. Data shown as mean ± 95% CI. Statistics by one-way ANOVA with Tukey’s multiple comparisons test (B,D,F,H) or Kruskal-Wallis with Dunn’s multiple comparisons test (C,F). See also Figure S3.

### Guar gum supplementation inhibits CD4^+^ Th1 activation in EAE

Autoreactive, encephalitogenic T cells are primed in secondary lymphoid organs prior to migration to the CNS in the MOG_35-55_ immunization model of EAE. Consistent with this, at d9 post-EAE induction during disease initiation, spleen size and total cellularity was increased in mice fed control and pectin-enriched diets. In contrast, splenic expansion was not seen in guar gum-fed mice following EAE induction (Figure 4A-B). The total number of activated CD44^hi^ CD4^+^ T cells was increased in control and pectin-fed spleens post-EAE induction, but was not apparent in guar gum-fed mice (Figure 4C-D).

**Figure 4.**
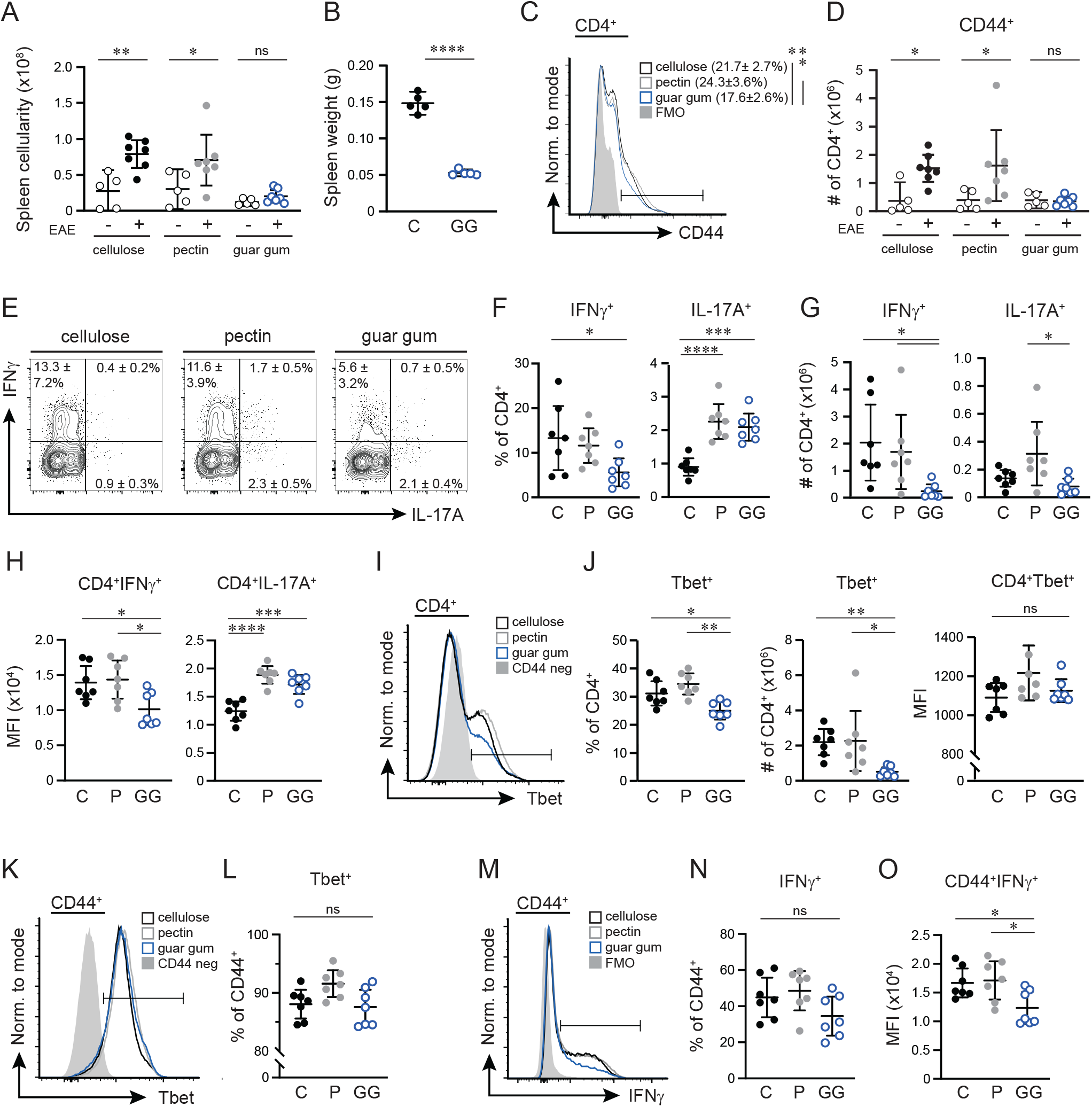
Guar gum supplementation inhibits CD4^+^ Th1 activation in EAE. Splenocytes were isolated at d9 post-EAE. **A)** Total spleen cellularity in naïve and EAE mice fed cellulose (C), pectin (P), or guar gum (GG). **B)** Spleen weights. **C,D)** CD4^+^ T cell activation assessed by CD44 expression. Concatenated histogram from one representative experiment. Proportion of CD44^+^ CD4 T cells shown as mean ± SD (C). Number of CD44-expressing CD4^+^ T cells in naïve and EAE mice (D). **E-H)** CD4^+^ cytokine expression following PMA/ionomycin stimulation. Representative, concatenated flow plot showing IFNγ and IL-17A expression in live CD45^+^ TCRβ^+^CD4^+^ cells, with mean ± SD of each group included (E). Proportion (F) and number (G) of CD4^+^ T cells expressing IFNγ or IL-17A. Median fluorescent intensity (MFI) of IFNγ and IL-17A of CD4^+^ T cells (H). **I-J)** Tbet expression in CD4^+^ T cells. Concatenated histogram of Tbet expression. Negative sample is CD44^-^ cells (I). Quantification of the proportion, number, and MFI of Tbet-expressing CD4^+^ T cells (J). **K-O)** Th1 polarization in CD44^hi^ cells. Concatenated histogram of Tbet expression in CD44^+^CD4^+^ T cells. Negative sample is CD44^-^ cells (K), quantified in (L). Concatenated histogram of IFNγ expression in CD44^+^CD4^+^ T cells. Negative sample is IFNγ FMO (M). Proportion (N) and MFI (O) of IFNγ^+^ CD44^+^CD4^+^T cells. Data shown as mean ± 95% CI. Statistics by 2-way ANOVA with Šídák’s multiple comparisons test (A,D), unpaired t-test (B), one-way ANOVA with Tukey’s multiple comparisons test (C,F,H,J,L,N,O) or Kruskal-Wallis with Dunn’s multiple comparisons test (G,J);

The diminished infiltration of Th1 cells into the CNS following MOG35-55 immunization may be due to impaired T cell activation, Th1 polarization, or both. To assess these possibilities, we quantified the total number of Th1 and Th17 cells in the spleen of control-, pectin-, and guar gum-fed mice. Interestingly, there was a small but significant increase in the frequency of IL-17A-producing CD4^+^ T cells in pectin- and guar gum-fed mice, and enhanced IL-17A expression on a per-cell basis (MFI) compared to control-fed mice (Figure 4E-H), but the numbers of Th17 cells were similar across all diets (Figure 4G). However, the frequency and total number of IFNγ^+^ CD4^+^ T cells, and per-cell expression of IFNγ, were all significantly reduced in guar gum-fed mice compared to control or pectin-fed mice (Figure 4E-H). Further, there were fewer CD4^+^ T cells expressing Tbet, the master transcription factor of Th1 cells in guar gum-fed mice, although per-cell expression remained similar across diets (Figure 4I-J).

These data suggest possible generalized defects in both T cell activation (determined by CD44 up-regulation) and Th1 polarization in CD4^+^ T cells of guar gum-fed mice. To determine whether Th1 polarization was impaired following activation, we measured Tbet and IFNγ expression within the small population of antigen-experienced CD44^hi^ CD4^+^ T cells. This analysis abrogated any differences in the proportions of Tbet^+^ or IFNγ^+^ cells in the CD44^hi^ population (Figure 4K-N), indicating that if cells can be initially activated in response to antigen exposure, they can acquire a Th1 phenotype. However, the IFNγ MFI remained lower in CD44^hi^ CD4^+^ T cells of guar gum-fed mice (Figure 4O), indicative of impaired effector function. Taken together, these data reveal a previously unappreciated role for the dietary fiber guar gum as a selective modifier of CD4^+^ T cell activation and Th1, but not Th17, polarization that interferes with the production of IFNγ.

### Impaired CD4^+^ T cell activation and Th1 polarization in guar gum-fed mice is cell intrinsic

Selective impairment of T cell activation and the Th1 differentiation program could be a consequence of T cell-intrinsic effects or external factors, such as altered dendritic cell (DC) activation, antigen presentation, or production of Th1-polarizing cytokines. To test this, CD4^+^ T cells from the spleens of control and guar gum-fed mice were cultured in the presence of a-CD3/a-CD28-coated beads under neutral (Thø), Th1, or Th17-skewing conditions. Under neutral conditions, the proliferative capacity of guar gum-derived CD4^+^ T cells was reduced compared to cellulose-derived CD4^+^ T cells, with a higher proportion and number of guar gum cells remaining undivided, and fewer cells reaching later stages of cell division (Figure 5A-B). Accordingly, the proliferation index (Figure 5C), the proportion of divided cells, the frequency of cells expressing Ki67, and the proportion of cells expressing CD44 (Figure S4A-D) were all diminished in guar gum-derived CD4^+^ T cells. Finally, although there were comparable concentrations of IL-17A in the culture media after 4 days of stimulation, guar gum-derived CD4^+^ T cells produced significantly less IFNγ than cellulose-derived cells (Figure 5D).

**Figure 5.**
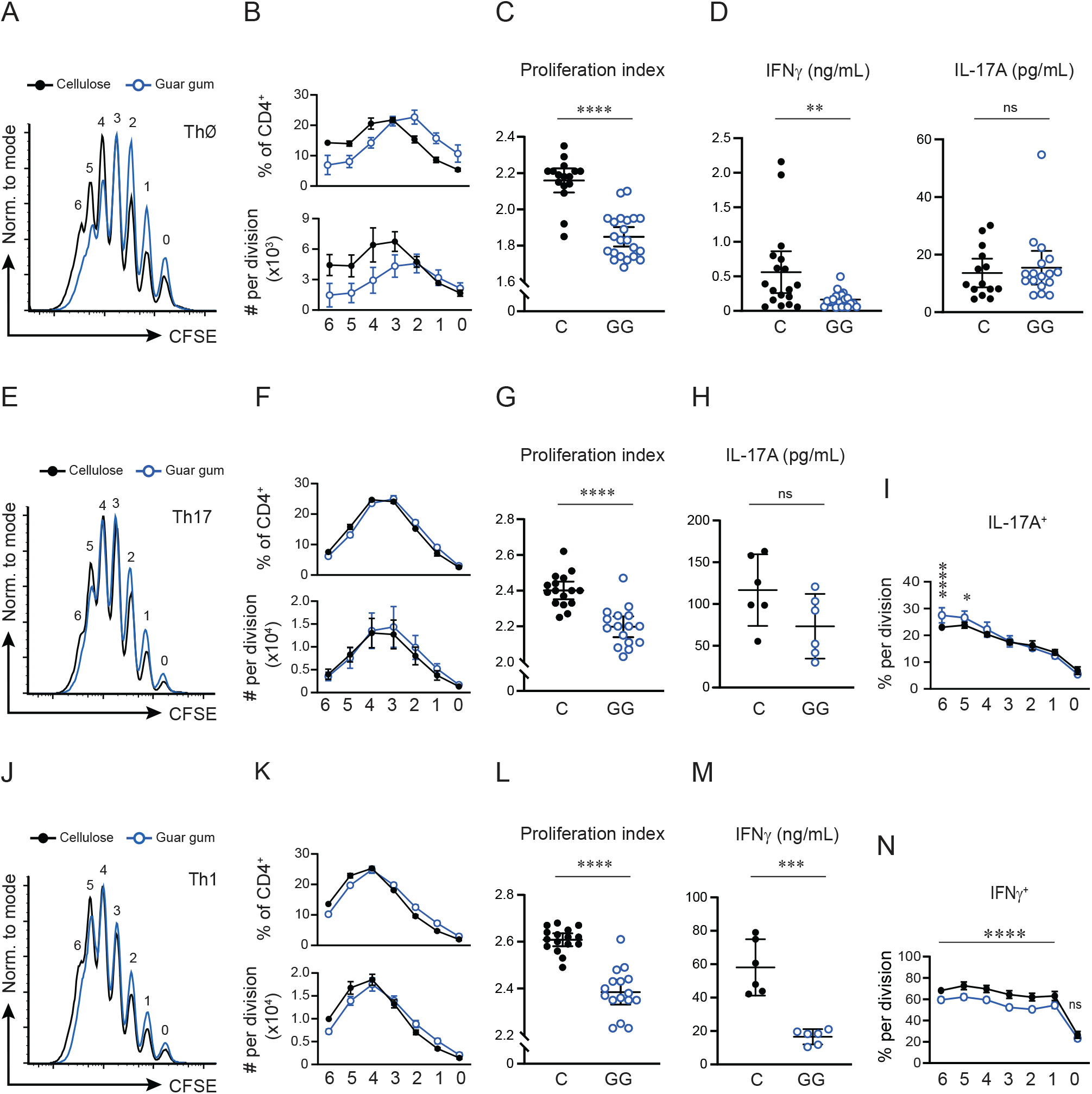
Reduced activation and polarization towards Th1 in guar gum cells is T cell-intrinsic. CD4^+^ T cells isolated from spleens of naïve cellulose (C)- or guar gum (GG)-fed mice, stained with CFSE, and cultured with a-CD3/a-CD28-coated beads for 96 hours in complete tissue culture media (Thø) **(A-D)**, in Th17-skewing conditions **(E-I),** or in Th1-skewing conditions **(J-N).** **A, E, J)** Representative histograms of CFSE dilution. **B, F, K)** Quantification of proliferation by the proportion (top) and number (bottom) of CD4^+^ T cells in each division. **C, G, L)** Proliferation index (total number of divisions / cells that went into division) of CD4^+^ T cells, calculated in FlowJo. **D, H, M)** Quantification of noted effector cytokines in the culture media after 96 hours by quantitative ELISA. **I, N)** Proportion of CD4^+^ T cells expressing noted cytokines at each stage of cell division. Data shown as mean ± 95% CI. Stats by Mann-Whitney test (C-D, G-H, L-M) or 2-way ANOVA with Šídák’s multiple comparisons test (I,N). See also Figure S4-5.

Under Th17-polarizing conditions, proliferation differences between guar gum- and control diet-derived cells were less apparent, although the statistically significant reduction in proliferation index and Ki67 expression by guar gum-derived cells was maintained (Figure 5E-G, Figure S4E-F). Consistent with Thø conditions, there was no difference in IL-17A secretion between cellulose- and guar gum-derived CD4^+^T cells, IL-17A expression in the bulk CD4^+^ population, or at each stage of cell division (Figure 5H-I, Figure S4G-H).

Consistent with the Thø and Th17-skewed conditions, guar gum-derived CD4^+^ T cells cultured under Th1 skewing conditions had a slight but statistically significant proliferative defect, measured by proliferation index, percent divided cells, and Ki67 detection (Figure 5J-L, Figure S4I-J). Under Th17 conditions, a similar effect had no impact on the overall amount of IL-17A produced. In contrast, there was a striking reduction in the concentration of IFNγ released into the culture supernatant of cells isolated from guar gum-fed mice (Figure 5M), which is consistent with the reduced frequency of IFNγ-producing cells at every stage of cell division (Figure 5N), and an overall diminution of the amount of IFNγ produced on a per-cell basis in the bulk population of guar gum-derived CD4^+^ T cells (Figure S4K-L). These data indicate that the reduced accumulation of IFNγ-producing CD4^+^ T cells in the CNS following EAE induction in guar gum-fed mice is at least partially due to a CD4^+^ T cell-intrinsic effect within the Th1, but not Th17, differentiation pathway. Further, these T cell-intrinsic activation defects are unique to guar gum and not a universal feature of high-fiber diets, as pectin-derived CD4^+^ T cells display similar proliferation and IFNγ production as cellulose-derived cells (Figure S5). Although these results do not rule out a contribution of DC-mediated impairment of T cell activation, they suggest that cell intrinsic differences are sufficient to account for the reduced IFNγ production seen *in vivo*.

### Guar gum supplementation is associated with changes in expression of migration and adhesion-related genes, including those necessary to cross the BBB

Some aspects of the ameliorated EAE phenotype cannot be explained simply by reduced IFNγ production in guar gum-fed mice, including the sweeping inability of T cells to migrate to the CNS. To expand our understanding of the changes occurring in the peripheral immune response following EAE induction, we designed a custom Nanostring nCounter panel to evaluate gene expression of several T cell activation, differentiation, and functional pathways in sort-purified CD4^+^ T cells isolated from cellulose- and guar gum-fed mice at d9 post-EAE (Figure 6A-B, Table S4).

**Figure 6.**
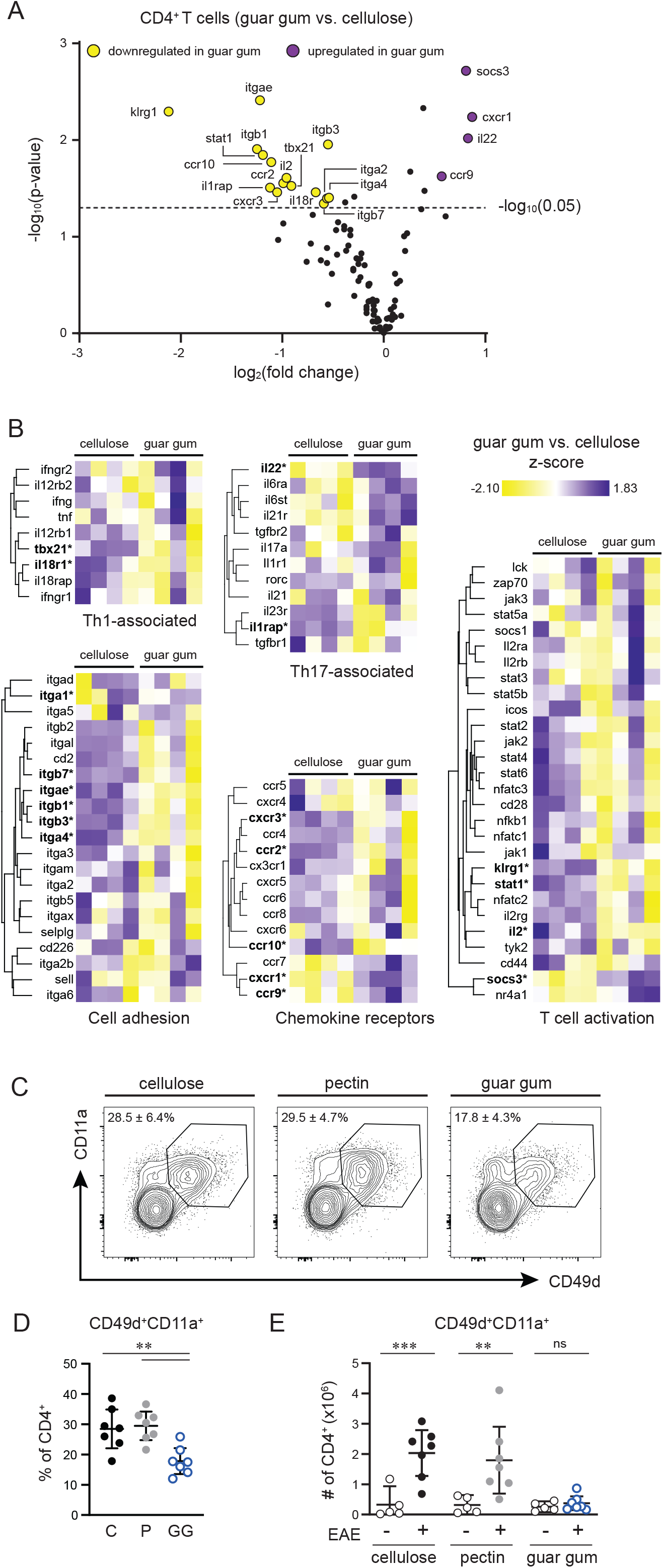
Guar gum supplementation associated with changes in expression of migration and adhesion-related genes, including those necessary to cross the blood-brain barrier. **A)** Differential gene expression from sort-purified CD4^+^ T cells isolated from control or guar gum-fed mice at d9 EAE by Nanostring nCounter (n=4 per group). Significance threshold: *p*-value = 0.05, log2(fold change) threshold = 0.5. **B)** Hierarchical clustering of gene expression for individual genes shown for individual mice. Significantly different genes are in bold with an asterisk (*). **C-E)** CD49d^+^CD11a^+^ expression of CD4^+^ T cells. Concatenated flow plot of from one representative experiment d9 post-EAE, with mean ± SD (C). Proportion (D) and number (E) of CD49d^+^CD11a^+^ CD4^+^ T cells. Data shown as mean ± 95% CI (D-E). Stats by one-way ANOVA (D) or two-way ANOVA with Šídák’s multiple comparisons test (E).

Consistent with protein-level expression, expression of *tbx21*, the gene encoding Tbet, was significantly reduced in EAE-induced guar gum-derived (guar gum/EAE) splenic CD4^+^ T cells. While changes in RNA-level expression of IFNγ or the IFNγ receptor were not detected, key regulators of IFNγ signaling (*socs3* and *stat1*) were up- and down-regulated, respectively, in guar gum/EAE CD4^+^ T cells. Additionally, expression of several other genes involved in IFNγwere diminished in guar gum/EAE CD4^+^ T cells, including IL-18Ra (*il18r1*) and IL-1 receptor accessory protein (IL1RAP, *il1rap*). IL-18/IL-18R interactions are centrally involved in Th1 polarization, and a lack of IL-18R has been shown to reduce T cell proliferation and limit IFNγ production and CD44 up-regulation in CD4^+^ T cells (Kaplanski, 2018; Oliveira et al., 2017). Further, formation of an IL-1RAP/IL-18Rα complex in response to IL-18 promotes IFNγ expression *in vitro* (Cheung H 2005 J Immunol).

Gene expression of *il2* and *klrg1* (another marker of CD4^+^ T cell antigen experience) were significantly down-regulated in guar gum/EAE cells, indicating an overall impairment in CD4^+^ T cell activation following EAE induction. Consistent with our observations of decreased Tbet expression and cytokine production, KLRG1 expression is directly downstream of Tbet signalling (Joshi et al., 2007), and KLRG1^+^ CD4^+^ T cells are potent cytokine-producers (Reiley et al., 2010). Further, *nr4a1* (Nur77) expression was trending upward in guar gum/EAE CD4^+^ T cells (log_2_(Fold Change): 0.61, *p*-value: 0.06). Nur77 is a faithful indicator of TCR activation, suggesting that immediate/early TCR signaling is not impaired by guar gum feeding. However, Nur77 acts as a negative regulator of EAE (Liebmann et al., 2018; Shaked et al., 2015; Wang et al., 2018), at least in part by acting as a “brake” on cellular metabolism to limit T cell proliferation (Liebmann et al., 2018).

Consistent with the inability of CD4^+^ T cells to infiltrate the CNS of guar gum-fed mice, gene expression of migration-associated receptors was widely affected in guar gum/EAE CD4^+^ T cells. The chemokine receptors *ccr10, ccr2, and cxcr3* were downregulated in guar gum/EAE cells, alongside upregulation of the gut homing-associated chemokine receptor genes *ccr9* and *cxcr1*. In MS patients, CCR9 has previously been demonstrated to be reduced in PBMCs (Kadowaki et al., 2019), and CXCR3^+^CD4^+^ T cells accumulate in CNS lesions and in the CSF (Sørensen et al., 1999; Sorensen et al., 2002). Further, CXCR3 is recognized as a marker of activation in Th1 cells due to direct activation of CXCR3 by Tbet (Lord et al., 2005) and is crucial for infiltration of Th1s into inflamed tissue (Qin et al., 1998).

Leukocyte-expressed integrins play an important role in the ability of cells to enter inflamed tissue by interacting with endothelial-expressed adhesion molecules. The integrin subunit genes *itga2* (CD49b), *itga4* (CD49d), *itgae* (CD103), *itgb1* (CD29), *itgb3* (CD61), and *itgb7* were expressed at significantly lower levels in guar gum/EAE CD4^+^ T cells. These alterations in migration-associated genes may be indicative of multi-faceted defects in the ability of CD4^+^ T cells to traffic to a site of inflammation and migrate into inflamed tissue, including across the blood-brain-barrier (BBB). Cellular migration across the BBB is mediated by the interactions of VLA-4 with VCAM-1, and LFA-1 with ICAM-1. Co-expression of VLA-4 (composed of integrins α4/CD49d and β1) and LFA-1 (composed of integrins αL/CD11a and β2), and thus the ability to migrate into inflamed tissue, can be used as an indicator of T cell activation and antigen experience (McDermott and Varga, 2011). Consistent with reduced expression of *itga4* (CD49d) and trending reduction in *itgal* (CD11a) (log_2_(Fold Change): −0.39, *p*-value: 0.07), the frequency of activated CD49d^+^CD11a^+^ CD4^+^ T cells with the capacity to cross the BBB was reduced in guar gum-fed mice compared to controls at d9 post-EAE induction (Figure 6C-D), and there was no detectable expansion of this population in guar gum-fed mice during EAE (Figure 6E).

### Defective Th1 activation and polarization is the primary driver of protection from EAE

Although our data indicate significant impairment of systemic Th1 activation and expression of migratory markers that may contribute to their impaired CNS infiltration, it is possible that other non-hematopoietic factors, such as BBB integrity and/or glial cell activation, could be altered following guar gum supplementation. Thus, we turned to an adoptive transfer model of EAE where the impact of guar gum supplementation on leukocyte activation can be separated from the effects on non-hematopoietic factors (Figure 7A).

**Figure 7.**
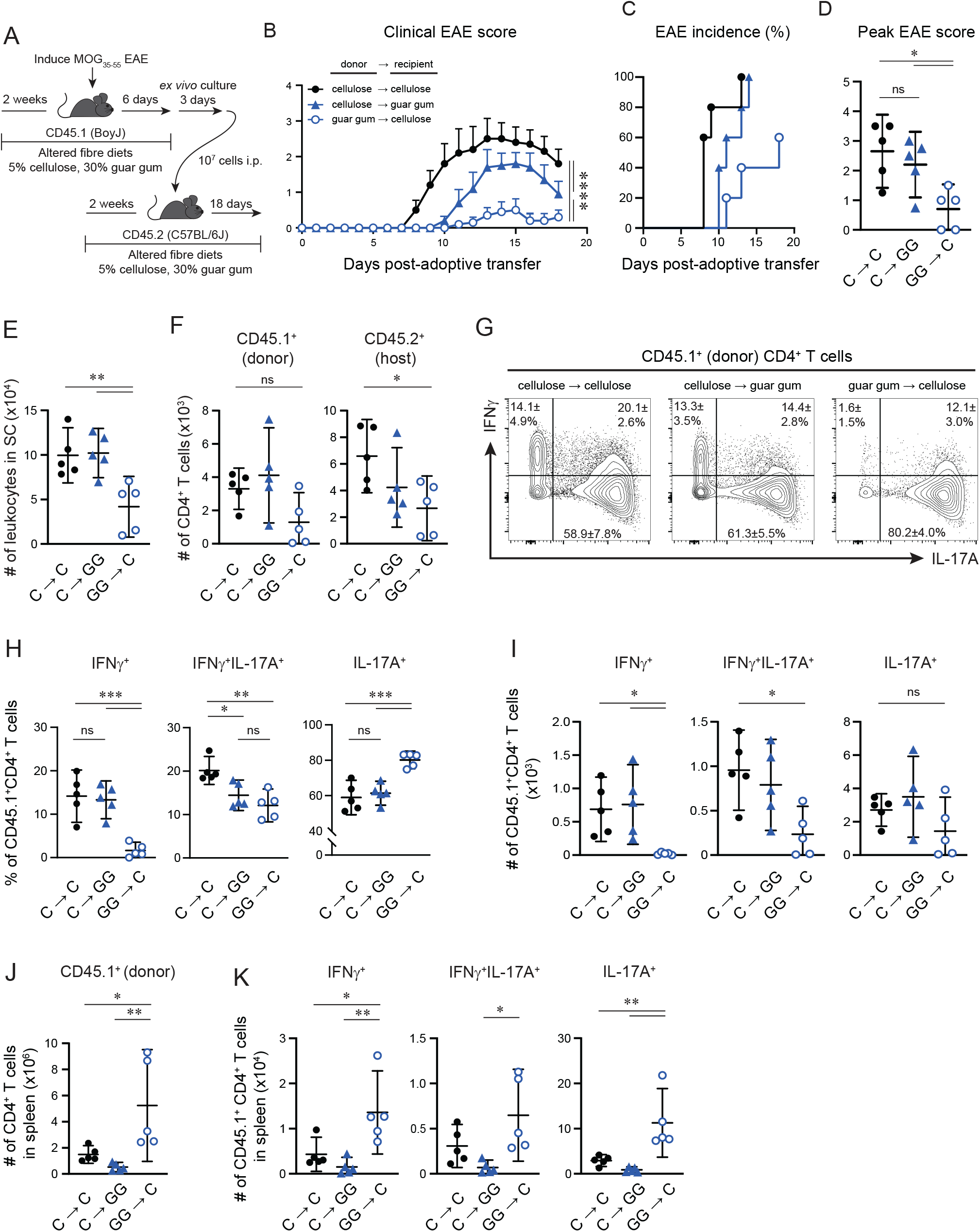
Defective Th1 activation and polarization is the primary driver of protection from EAE. **A)** CD45.1 ^+^ mice are fed a 5% cellulose (C) or 30% guar gum (GG) diet for 2 weeks prior to MOG35-55 immunization. Encephalitogenic donor cells are collected, cultured, and injected i.p. into recipient C57BL/6 (CD45.2^+^) mice fed with 5% cellulose or 30% guar gum for 2 weeks prior to adoptive transfer. Cellulose donor cells transferred to cellulose-fed recipients (C→C); cellulose donor cells transferred to guar gum-fed recipients (C→GG); guar gum donor cells transferred to cellulose-fed recipients (GG→C). **B-D)** Outcomes of mice that received adoptive transfers: daily clinical scores (B), incidence (C), and peak score (D). **E)** Total number of leukocytes in the spinal cord (SC) by flow cytometry forward and side scatter. **F)** Number of CD45.1^+^ (donor) and CD45.2^+^ (host) CD4^+^ T cells in the SC day 18 post-EAE. **G-I)** IFNγ and IL-17A expression of CD45.1^+^ donor cells recovered from SC of recipient mice. Concatenated flow plots showing mean ± SD (G). Proportion (H) and number (I) of IFNγ^+^, IFNγ^+^IL-17A^+^, and IL-17A^+^ CD45.1^+^CD4^+^ T cells. **J)** Number of CD45.1^+^ (donor) CD4^+^ T cells in the spleen d18 post-EAE. **K)** Number of IFNγ^+^, IFNγ^+^IL-17A^+^, and IL-17A^+^ CD45.1^+^CD4^+^ T cells in the spleen. Data shown as mean ± 95%CI. Stats by two-way ANOVA with Šídák’s multiple comparison test (column factor shown for each comparison), one-way ANOVA with Holm-Šídák multiple comparisons test (D-F, H-K).

Transfer of CD45.1 ^+^ donor cells isolated from control-fed mice into control-fed recipients (C→C) reliably induced ascending paralysis (median symptom onset (d): 8, range: 8-13). Interestingly, transfer of control-fed cells to guar gum-fed recipients (C→GG) slightly delayed disease onset (median symptom onset (d): 11, range 10-14) (Figure 7B-C). Nevertheless, both cohorts that received CD45.1 ^+^ cells from a cellulose-fed donor developed similar disease incidence and severity by d18 post adoptive transfer (Figure 7C-D), suggesting that the encephalitogenic capacity of the cellulose donor cells is sufficient to overcome any non-hematopoietic protective factors provided by the guar gum diet. In contrast, transfer of an equivalent number of guar gum-derived donor cells into control-fed recipients (GG→C) dramatically altered clinical outcomes, with reduced incidence and disease severity (Figure 7B-D). The encephalitogenic T cells activated in a guar gum-exposed environment, therefore, are insufficient to induce severe neuroinflammation in a control-fed host.

We assessed the differentiation and localization of the adoptively transferred cells at d18 post-adoptive transfer. Mice that received encephalitogenic cells from control-fed donor mice had similar numbers of total infiltrating leukocytes in the SC regardless of host diet (C→C and C→GG) (Figure 7E). In contrast, leukocyte infiltration of the SC was significantly reduced in GG→C mice, which was primarily driven by reduced recruitment of host, rather than donor, cells (Figure 7E-F). Notably, although there was no significant difference in the overall number of donor CD4^+^ T cells reaching the SC across all groups, there were significantly fewer IFNγ^+^ and IL-17A^+^IFNγ^+^ double-producing, but not IL-17A^+^ donor cells isolated from GG→C mice (Figure 7G-I). This blunting of the CNS-infiltrating Th1 population, but not of Th17s, is reminiscent of the phenotype observed during active EAE and highlights the importance of the Th1 compartment in mediating neuroinflammation in this model.

Finally, supporting the hypothesis that cellular migration is altered in guar gum/EAE cells, CD45.1^+^ donor cells were significantly more abundant in the spleen of GG→C mice compared to either the C→C or C→GG cohorts (Figure 7J), indicating disrupted migration of guar gum-derived donor cells to the site of inflammation. Notably, IFNγ^+^, IFNγ^+^IL-17A^+^ double-positive, and IL-17A^+^ CD4^+^ T cells were all increased in the spleens of GG→C mice compared to the cellulose-derived cells (Figure 7K). Collectively, these data provide critical insight into the physiological effects of dietary fiber supplementation, demonstrating that distinct fiber types can tune specific elements of immunoreactivity. Deeper understanding of the effects of different fiber types on the host could lead to more effective deployment of dietary interventions in disease states.

## Discussion

The association between microbiota-mediated metabolism of dietary fibers and the downstream effects on host health and immune function have been firmly established in both humans and animal models (Barber et al., 2020; Cai et al., 2020). Invariably, fiber supplementation induces shifts in microbial community composition and function, with a common outcome of increased SCFA availability. The effects of fiber supplementation on host physiology include enhanced numbers and function of CD4^+^ Treg cells, support of cellular metabolism, and (with few exceptions (Komatsu et al., 2021; Myhill et al., 2020; Prow et al., 2019) restrained inflammatory responses that protect against infectious and non-communicable inflammatory disease models in mice. Some of the studies leading to these insights have analyzed supplementation with single-source high fiber diets (pectin, inulin, cellulose) while others have used mixed fiber sources. As the field of host-microbiome-diet cooperativity develops, we need a more granular understanding of how different fiber types are used by the microbiota and the overall effect on host physiology and immune function. To this end, we compared different types of single-source soluble fiber (pectin, inulin, resistant starch, guar gum) and demonstrate a previously undescribed signature of modified Th1 activation in response to one source (guar gum) that limits the accumulation of pathogenic inflammatory cells in the CNS and ameliorates EAE.

Guar gum is a large plant-based soluble fiber composed of repeating units of galactose side chains on a mannopyranose backbone, commonly used as an emulsifier or thickener in processed foods (Mudgil et al., 2014). Supplementation with guar gum or a derivative (partially hydrolyzed guar gum) in small animal models and humans can reduce cholesterol and other blood lipids (Moriceau et al., 2000; Superko et al., 1988; Yasukawa et al., 2012), and has been studied in the context of inflammatory bowel diseases as a means of supporting epithelial barrier integrity and wound healing (Horii et al., 2016; van Hung and Suzuki, 2016; Naito et al., 2006; Takagi et al., 2016). However, the effects of guar gum on immune cell function have been understudied. Although guar gum supplementation, much like other soluble dietary fibers, enhanced cecal SCFA levels, it is unlikely that SCFA accounts for the potent anti-inflammatory properties that protect mice from neuroinflammation. Despite clear differences in clinical outcomes between pectin- and guar gum-supplemented mice, the profiles of SCFA availability in these cohorts of mice were quite similar. SCFA supplementation in EAE is associated with increased numbers of IL-10-, IL-17A-, and IFNγ-producing CD4^+^ T cells, likely via increased glycolytic capacity of activated leukocytes (Park et al., 2019). Our data suggests that guar gum feeding impairs CD4^+^ T cell IFNγ production *in vivo* and *in vitro*, even under strong Th1 polarization conditions. Impaired Th1 polarization and the associated migratory defects suggested by gene expression analysis and *in vivo* adoptive transfers have not previously been associated with SCFA supplementation, and our data suggest that guar gum supplementation triggers an alternative mechanism of T cell immunomodulation that may be SCFA-independent.

Microbiota-mediated breakdown of dietary components, and the accompanying changes to microbiota metabolism, release an abundance of small metabolites beyond SCFAs that can have potent immunomodulatory functions, including bile acids and tryptophan-derived molecules that have been implicated in regulation of the gut-brain axis and MS (Bhargava et al., 2020; Rothhammer et al., 2018). Although the identity of the specific metabolite(s) that are responsible for impaired Th1 activation in guar gum-fed mice remain unidentified, future work specifically addressing the microbial ecological networks that respond to guar gum and other dietary fiber sources to produce different suites of immunomodulatory compounds could be transformative for our understanding of gut-to-brain signaling and personalized dietary recommendations.

Importantly, although the MOG_35-55_ EAE model in C57BL/6 mice is generally associated with Th17-mediated pathology, the data presented here align with a model of collaboration between myelin-reactive Th1 and Th17 cells. IFNγ may have a particularly important role in initiation phase of EAE. Supporting this, myelin-specific 2D2 TCR transgenic cells primed in the absence of pertussis toxin do not produce IFNγ or induce disease, despite similar IL-17A expression compared to pertussis toxin-primed controls (Ronchi et al., 2016). Furthermore, induction of EAE by adoptive transfer of Th17 polarized cells in C57BL/6 mice is concurrent with appearance of IFNγ-producing cells (O’Connor et al., 2008), which may be due to IFNγ-mediated upregulation of the adhesion molecules ICAM and VCAM on CNS endothelial cells that facilitates interaction with leukocyte expressed integrins and transmigration of inflammatory CD4^+^ T cells across the BBB (Loos et al., 2020).

Collectively, the data presented here advance our understanding of diet-microbiota-immune cross-talk and identifies a new role for a particular fermentable fiber source, guar gum, in the modulation of CD4^+^Th1 activation and differentiation that confers protection from autoimmune-mediated paralysis. Continued exploration of fiber degradation pathways of specific microbes, the metabolites they produce, and the signaling programs induced in leukocytes is needed to effectively harness the immunomodulatory potential of dietary interventions.

## Materials & Methods

### Mice

Mice were housed at the University of British Columbia under specific pathogen-free conditions at the Center for Disease Modeling or the Modified Barrier Facility. C57BL/6J (000664) and BoyJ (B6.SJL-*Ptprc*^a^*Pepc*^b^/BoyJ, 002014) mice were ordered from The Jackson Laboratories or bred in the facility. If ordered in, 4-6 week old female mice were randomized into new cages within 48 hours of delivery to minimize cage-specific effects, and allowed to acclimatize for at least one week before exposure to experimental diets. Experimental mice were age- and sex-matched within each experiment and were between 6-12 weeks of age for all experiments. All experiments were performed according to guidelines from UBC Animal Care Committee and Biosafety Committee-approved protocols.

In the Centre for Disease Modeling, up to 5 mice per cage were housed in ventilated Ehret cages prepared with BetaChip bedding and had *ad libitum* access to experimental diets and reverse osmosis/chlorinated (2-3 ppm)-purified water. Housing rooms were maintained on a 14/10-hour light/dark cycle with temperate and humidity ranges of 20-22°C and 40-70%, respectively. Sentinel mice housed in experimental rooms were maintained on dirty bedding and nesting material and were tested on a quarterly basis for presence of mites (*Myocoptes, Radford/Myovia*), pinworm (*Aspiculuris tetaptera, Syphacia obvelata*), fungi (*encephalitozooan cuniculi*), bacteria (*Helicobacter spp., Clostridium piliforme, Mycoplasma pulmonis*, CAR Bacillus), and viruses (Ectromelia, EDIM/Rotavirus, MHV, MNV, MVM, LCMV, MAV1/2, MCMV, Polyoma, PVM, REO3, Sendai, and TMEV).

In the Modified Barrier Facility, up to 5 mice per cage were housed in OptiMouse cages prepared with BetaChip bedding and had *ad libitum* access to experimental diets and reverse osmosis/chlorinated (2-3 ppm)-purified water provided in sealed polycarbonate bags accessible via sterile sipping valves. Housing rooms were maintained on a 14.5-9.5-hour light/dark cycle with temperature and humidity ranges of 22-25°C and 50-70%, respectively. Sentinel mice housed in experimental rooms were maintained on dirty bedding and nesting material and were tested on a quarterly basis for presence of parasites (pinworms and fur/follicular mites, *Pneumocystis spp*. (*carinii, murina*)), bacteria (*Mycoplasma pulmonis*), and viruses (RADIL Comprehensive Panel) including Mouse Hepatitis Virus (MHV) Mouse Minute Virus (MMV), Mouse Parvovirus (NS1 – Generic Parvovirus & MPV 1-5), Theiler’s Murine Encephalitis Virus (TMEV), Epizootic Diarrhea of Infant Mice (EDIM), Sendai Virus, Pneumonia Virus of Mice (PVM), Reo3 virus (REO3), Lymphocytic Choriomeningitis Virus (LCMV), Ectromelia virus, Murine Adenovirus I/II (MAV1/MAV11), Polyomavirus.

### Diets

Mice were given *ad libitum* access to an experimental diet for 2 weeks prior to any given experiment and maintained on the same diet over the course of any experimental procedures. Altered-fiber diets were custom-ordered from TestDiet and built upon a common standard diet (AIN-93G, 57W5), and thus contained similar levels of vitamins, minerals, fat, and protein. Full dietary information and ingredients can be found in Supplemental Table 1. For all experiments, a control diet of 5% cellulose (57W5), the base diet for all other modified diets, was included. Experimental diets include: 5% cellulose (AIN-93G-57W5), 0%fiber (AIN-93G-9GKZ), 30% guar gum (AIN-93G-5BSE), 30% pectin (AIN-93G-5BSX), 30% inulin (AIN-93G-5BX1), and 30% resistant starch (AIN-93G-5BAC).

### EAE induction

EAE was induced by immunization of 8-12 week-old mice with 200 μg of MOG_35-55_ peptide (GenScript) emulsified in complete Freund’s Adjuvant (400 μg desiccated *Mycobacterium tuberculosis* H37 Ra mixed with incomplete Freund’s adjuvant (BD Biosciences)). The emulsion was injected subcutaneously in a total volume of 100 μL in the hind flank under isoflurane anaesthesia. Intraperitoneal (i.p.) injections of 200 ng pertussis toxin (List Biological Laboratories) were administered on the day of immunization and 48 hours later.

Adoptive transfer EAE was induced by immunization of donor mice as described above, with pertussis toxin administered on the day of immunization only. Spleens, and inguinal and brachial lymph nodes were collected day 6 post-EAE induction, processed into single cell suspensions, and cultured at 37°C in RPMI-based culture media (10% FBS, L-glutamine, Penicillin/streptomycin, 1X MEM NEAA (Sigma), 1mM sodium pyruvate (Sigma), 100 mM HEPES, 55 μM β-mercaptoethanol) supplemented with 20 μg/mL MOG_35-55_ peptide (GenScript), 20 ng/mL IL-12 (Biolegend), 20 ng/mL IL-23 (R&D Biosystems), 20 ng/mL IL-6, (Biolegend) and 4 ng/mL TGF-β (Biolegend) for mixed Th1/Th17 skewing conditions. After 72 hrs of culture, 10^7^ live leukocytes were i.p. injected into naïve recipient mice.

Mice were evaluated for clinical health and EAE severity daily after the onset of EAE symptoms. EAE severity was scored on a 5-point scale: 0, no clinical symptoms; 0.5, partial loss of tail tonicity; 1, paralyzed tail; 1.5, loss of coordinated movement; 2, hindlimb paresis; 2.5, paralysis in one hind limb; 3, paralysis in both hind limbs; 3.5, both hindlimbs paralyzed accompanied by weakness in forelimbs; 4, paralysis in at least one forelimb (humane endpoint); 5, moribund or dead. Mice that died during the course of EAE were assigned a 5 on the day of death then removed from further analysis. Only mice that survived until dedicated experimental endpoints were induced in area under the curve calculations.

### Histology

Mice were euthanized by CO_2_ inhalation and transcardially perfused with at least 20 mL cold PBS on day 15 post-EAE. Brain and SCs were soaked in 10% neutral buffered formalin for 24 hours then transferred to 70% ethanol for storage at 4°C prior to paraffin embedding. Brains were cut down the midline for sagittal sectioning, and SCs were cut into 4 approximately equal-length sections for cross-sectional tissue slices at approximate levels of C2, C7, T8, L1. Paraffin blocks were sectioned at a thickness of 5 μm and serial sections were stained with hematoxylin and eosin (H&E) or Luxol Fast Blue (LFB). Paraffin embedding, sectioning, and staining were completed by a commercial histology service (Wax-it Histology Services Inc.). Slides were imaged with a Zeiss Axio Observer Z1 and an AxioCam 105 microscope camera. Brightfiled mages acquired using TL LED light intensity of 33.4% and 3.067ms exposure with a 10X objective. Images were processed in Adobe Photoshop to remove the background.

### Histology quantification

4 images per SC (one at each SC level described above) were scored for infiltration in a blinded manner by 2 investigators. The mean score between observers was used for quantification. Inflammation/infiltration scores were assigned using a 4-point system as follows: 0, no infiltration; 1, mild infiltration in perivascular space; 2, several separate regions of perivascular infiltration/cuffing; 3, severe perivascular infiltration and 1-2 regions of parenchymal infiltration; 4, severe diffuse infiltration with several parenchymal clusters. The cerebellum for each mouse was scored for infiltration in a similar blinded manner using a 4-point system: 0, no infiltration; 1, small clusters of cells into parenchymal space; 2, mild infiltration at 1-2 regions; 3, severe infiltration and 1 other region of mild infiltration; 4, severe infiltration at 2 or more regions.

### Leukocyte recovery and *ex vivo* stimulation

Single cell suspensions of splenic leukocytes were prepared by mechanical dissociation and passage through a 70 μm nylon mesh filter followed by ACK lysis to remove red blood cells. To isolate leukocytes from brain and SC tissue, mice were transcardially perfused with at least 20mL cold PBS, then tissues were passed through a 70 μm nylon mesh filter followed by resuspension in room temperature 40% isotonic Percoll and centrifugation at 500 ×g for 15 minutes with no brake. Cell pellets were resuspended in FACS buffer (PBS supplemented with 2% NCS and 1mM EDTA) or in stimulation media for intracellular staining for cytokine responses. For polyclonal cytokine responses, single cell suspensions were stimulated with 0.1 μg/mL PMA (Sigma) and 0.1 μg/mL ionomycin (Sigma) in the presence of Brefeldin A (eBioscience/ThermoFisher) and GolgiSTOP (BD Biosciences) for 5 hours at 37°C.

### Flow cytometry and Fluorescence-activated cell sorting

Cells were washed with cold PBS prior to staining with live/dead Fixable Aqua viability dye (Invitrogen/ThermoFisher) for 15 minutes at 4°C in the dark. Subsequently, cells were washed with FACS buffer and incubated with 2 μg/mL α-mFcγR (clone 24G2, AbLab) for 5 minutes at 4°C. Pre-determined concentrations of fluorophore-labelled antibodies were added for a final volume of 100uL of Fc-block and antibodies in FACS buffer. Wells were mixed thoroughly then incubated for 20 minutes at 4°C in the dark. If intracellular staining was being performed, cells were next incubated with FOXP3/Transcription Factor Fixation/Permeabilization buffer (eBioscience/ThermoFisher) for 30 minutes at 4°C in the dark, followed by 3 washes with permeabilization buffer. Fluorophore-labelled antibodies for intracellular targets were diluted in 100 μL permeabilization buffer per well and incubated with cells for 40 minutes at room temperature in the dark. The following antibodies were used for analysis of brain and spinal cord tissue: CD45.2-AlexaFluor700 (clone 104, Biolegend), TCRß-PerCP-Cy5.5 (H57-597, Biolegend), B220-APC-eFluor 780 (RA3-6B2, eBioscience), CD4-PE-Cy7 (GK1.5, eBioscience), CD8α-BV650 (53-6.7, BD), IFNγ-eFluor450 (XMG1.2, eBioscience), IL-17A-PE/Dazzle 594 (TC11-18H10.1, Biolegend), FOXP3-AlexaFluor488 (FJK-16S, eBioscience). The following antibodies were used for analysis of spleen tissue d9 post-EAE: CD45.1-AlexaFluor700 (A20, BioLegend) or CD45.1-PE (A20, BioLegend), CD45.2-AlexaFluor700 (104, BioLegend), TCRb-PE/Dazzle 594 (H57-597, BioLegend), B220-PE (RA3-6B2, BioLegend), CD4-BV650 (RM4-5, BD Bioscience), CD8a-PE/Cy7 (53-6.7, BD Bioscience), CD44-APC/Cy7 (IM7, BioLegend), IFNγ-APC (XMG1.2, eBioscience), IL-17A-eFluor450 (eBio17B7, eBioscience), FOXP3-AlexaFluor488 (FJK-16S, eBioscience), CD11a-PerCP-Cy5.5 (2D7, BD Bioscience), CD49d-FITC (R1-2, BioLegend), Tbet-APC (4B10, BioLegend), RORgt-PE (AFKJS-9, eBioscience).

Samples were run on LSRII-561 (BD Biosciences), Attune NxT (ThermoFisher), or CytoFLEX LX N3-V5-B3-Y5-R3-I0 (BD Biosciences) flow cytometers. Analysis was performed using FlowJo v10.7 (BD Life Sciences). FACS sorting was performed on an Influx-2 on pre-enriched CD4^+^ T cells (enrichment using an EasySep CD4^+^ T cell isolation kit (StemCell Technologies)). CD4^+^ T cells (C57BL/6J splenocytes). 10^6^ cells (CD45.2^+^TCRß^+^CD4^+^ and CD8a^-^B220^-^F4/80^-^CD11b^-^) were sorted into FBS-supplemented PBS on ice.

### Nanostring nCounter analysis

RNA was extracted using the PureLink RNA Mini Kit (Invitrogen) as per manufacturer’s instructions from 10^6^ FACS-sorted CD4^+^ T cells. RNA was analyzed for RNA integrity and concentration using an Agilent 22100 BioAnalyzer. 100 ng of RNA per sample was run on a custom Nanostring nCounter panel (see Table S4 for full gene list). Analysis was performed using nSolver2.0 (NanoString). Each sample passed hybridization quality controls. Transcripts were removed with numbers of reads below the background threshold determined by negative controls. Data was normalized to positive controls and to 4 housekeeping genes (*gusb, hprt, polr1b, rpl19*) which were not differentially expressed across samples (*p*>0.05). Statistical analysis in nSolver2.0 are t-tests. Distribution of t-statistics calculated using the Welch-Satterthwaite equation for degrees of freedom in the estimation of 95% confidence limits.

### Proliferation and Th1/Th17 skewing

Spleens were processed into single cell suspensions and CD4^+^ T cells were isolated using the EasySep CD4^+^ T cell isolation kit (StemCell Technologies) as per manufacturer’s instructions. Isolated CD4^+^ T cells (5×10^6^ cells/mL) were stained with 2.5 μM carboxyfluorescein succinimidyl ester (CFSE) in the dark at 4°C for 8 minutes prior to FCS quenching. CFSE-stained cells (10^5^ per well) were plated at a 1:1 ratio with Dynabeads™ mouse T-activator CD3/CD28 (Invitrogen) and cultured at 37°C in RPMI-based culture media for 96 hours. Supernatants were collected for cytokine profiling by quantitative ELISA for IFNγ and IL-17A as per manufacturer’s instructions (Invitrogen). Prior to flow cytometric analysis for cytokine production, cells were resuspended with 0.1 μg/mL PMA and 0.1 μg/mL ionomycin in the presence of brefeldin A and GolgiSTOP for 5 hours at 37°C.

For Th1 and Th17 skewing conditions, CD4^+^ T cells were isolated from the spleen and plated at a 1:1 ratio with αCD3/αCD28 beads as described above. For Th1 conditions, cells were cultured in the presence of 10 ng/mL IL-12, 10 ng/mL IL-2, and 10 μg/mL a-IL-4 (clone 11B11, BioXCell). For Th17 conditions, cells were cultured in the presence of 10 ng/mL IL-23, 10 ng/mL IL-2, 20 ng/mL IL-6, 5 ng/mL TGF-β1, 10 μg/mL aIL-4, and 10 μg/mL αIFNγ (clone XMG1.2, BioXCell). Cultures were analyzed for cytokine production by quantitative ELISA and flow cytometry as described above.

### Short chain fatty acid quantification

Major short chain fatty acids (SCFAs) were analyzed from cecal and serum samples as described previously (Brown K ISME 2016). Flash-frozen tissue samples were homogenized in isopropyl alcohol, containing 2-ethyl butyric acid at 0.01% v/v used as an internal standard and then centrifuged. The supernatant was injected into a Trace 1300 Gas Chromatograph, equipped with flame-ionization detector, with AI1310 autosampler (Thermo Scientific, Walkham, MA, USA) in splitless mode. A fused-silica FAMEWAX (Restekas, Bellefonte, PA, USA) column 30 m × 0.32 mm i.d. coated with 0.25 μm film thickness was used. Helium worked as the carrier gas at a flow rate of 1.8 ml/min. The initial oven temperature was 80°C, maintained for 5 min, raised to 90 °C at 5 °C/min, then increased to 105°C at 0.9 °C/min, and finally increased to 240°C at 20 °C/min and held for 5 min. The temperature of the flame-ionization detector and the injection port was 240°C and 230°C, respectively. The flow rates of hydrogen, air and nitrogen as makeup gas were 30, 300 and 20 mL/min, respectively. Data was analyzed with Chromeleon 7 software (Bannockburn, IL, USA). Volatile-free acid mix served as the standard to confirm the individual SCFAs (Sigma, Oakville, ON, Canada).

### Fecal DNA extraction, amplification, and iTag sequencing

DNA was extracted from fecal pellets using DNeasy PowerSoil Kit (Qiagen) as per manufacturer’s instructions. Resulting DNA was stored at −20°C. Purity and quality of DNA was assessed based on ratios for A_260_/A_280_ measured using a NanoDrop®ND-1000 spectrophotometer (Thermo Scientific). Bacterial and archaeal SSU rRNA gene fragments from the extracted genomic DNA were amplified using primers 515F and 806R. Sample preparation for amplicon sequencing was performed as described as in (Apprill et al., 2015; Caporaso et al., 2011). The amplicon library was analyzed on an Agilent Bioanalyzer using the High Sensitivity DS DNA assay to determine approximate library fragment size, and to verify library integrity. Pooled library concentration was determined using the KAPA Library Quantification Kit for Illumina. Library pools were diluted to 4 nM and denatured into single strands using fresh 0.2 N NaOH as recommended by Illumina. The final library was loaded at a concentration of 8 pM, with an additional PhiX spike-in of 5-20%. Sequencing was conducted at the UBC Sequencing and Bioinformatics Consortium (https://sequencing.ubc.ca/).

### Bioinformatic analysis

Sequences were processed using Mothur (Schloss et al., 2009). In brief, sequences were removed from the analysis if they contained ambiguous characters, had homopolymers longer than 8 bp and did not align to a reference alignment of the correct sequencing region. Unique sequences and their frequency in each sample were identified and then a pre-clustering algorithm was used to further de-noise sequences within each sample (Schloss et al., 2009). Unique sequences were identified and aligned against a SILVA alignment (available online at http://www.mothur.org/wiki/Silva_reference_alignment). Sequences were chimera checked using UCHIME (Edgar et al., 2011) and reads were then clustered into 97% OTUs using OptiClust (Westcott and Schloss, 2017). OTUs were classified using SILVA reference taxonomy database (release 138, available at http://www.mothur.org/wiki/Silva_reference_files). Singletons (OTUs with one 16s rRNA sequence read) were removed from analysis. All data was visualized in R and Excel. For alpha and beta diversity measures, all samples were subsampled to the lowest coverage depth (n=29256) and calculated in Mothur; community structure was investigated using the Yue and Clayton similarity estimator and a Principle Coordinates (PCoA) was used to compare bacteria community structures across all samples, and a Multiple sample Analysis of Molecular Variance (AMOVA) was used to test for significance of differences between microbial community populations. Non-metric multidimensional scaling of Bray-Curtis dissimilarity values based on OTU relative abundances was plotted using the vegan package in R. The highly significant correlations (R-squared greater than 0.5) between the ordination configuration and OTU abundances were identified using the envifit function in vegan and included as representative points on the NMDS plot.

To test for more nuanced microbial community responses, a statistical analysis, based on Linear discriminant analysis Effect Size (LEfSe) (Segata et al., 2011), was to identify microbial species that respond to different diets. Additionally, Indicator Value Analysis (IndVal) implemented in the R package indicspecies (de Cáceres and Legendre, 2009) was used to determine statistically significant associations between taxa and a treatment group, in this case, diet, using permutation tests. Indicator values are based on fidelity, or the probability of an OTU being found in all samples of a given group, and specificity, or the probability of the OTU being found in only one group given its presence in that group (Dufrene and Legendre 1997).

### Statistical Analysis

Statistical analyses were performed using GraphPad Prism (GraphPad Software, Version 9.0). Data was assessed for normal distribution by Shapiro-Wilk test to determine the appropriate statistical test. Two-tailed nonparametric Mann-Whitney test, two-tailed t-tests, one-way ANOVA with Tukey’s multiple comparisons test, Kruskal-Wallis with Dunn’s multiple comparisons test, 2-way ANOVA with Tukey’s multiple comparisons test or Šídák’s multiple comparisons test were used as specified in figure legends. Values are reported as mean ± 95% confidence interval unless otherwise stated. Data represented alongside representative flow plots depict mean ± SD. Symbols: ns p> 0.05, * p< 0.05; ** p<0.01; *** p<0.001; **** p<0.0001.

## Acknowledgments

Authors acknowledge support from the MS Society of Canada endMS Personnel Award program (NMF, JRA), the Canadian Institutes for Health Research (PJT-148909) and the Canada Research Chair program (LCO). We are grateful for excellent technical support from ubcFLOW, the Centre for Disease Modeling, the Modified Barrier Facility, the UBC Sequencing and Bioinformatics Consortium, the Centre for Heart and Lung Innovation Molecular Phenotyping Core.

## Contributions

NMF, HGR, and LCO conceived the study. NMF and HGR led experimental design, execution and analyzed data for *in vivo* and *in vitro* studies with support from JRA, NS, JHS, EJW. RLS and SAC ran analysis of microbial communities. YJ and DLG performed quantification of SCFA. NMF and LCO wrote the manuscript.

