## Supplemental data for "The dietary fiber guar gum ameliorates experimental autoimmune encephalomyelitis via attenuated Th1 activation and differentiation"

### Supplemental Figure 1

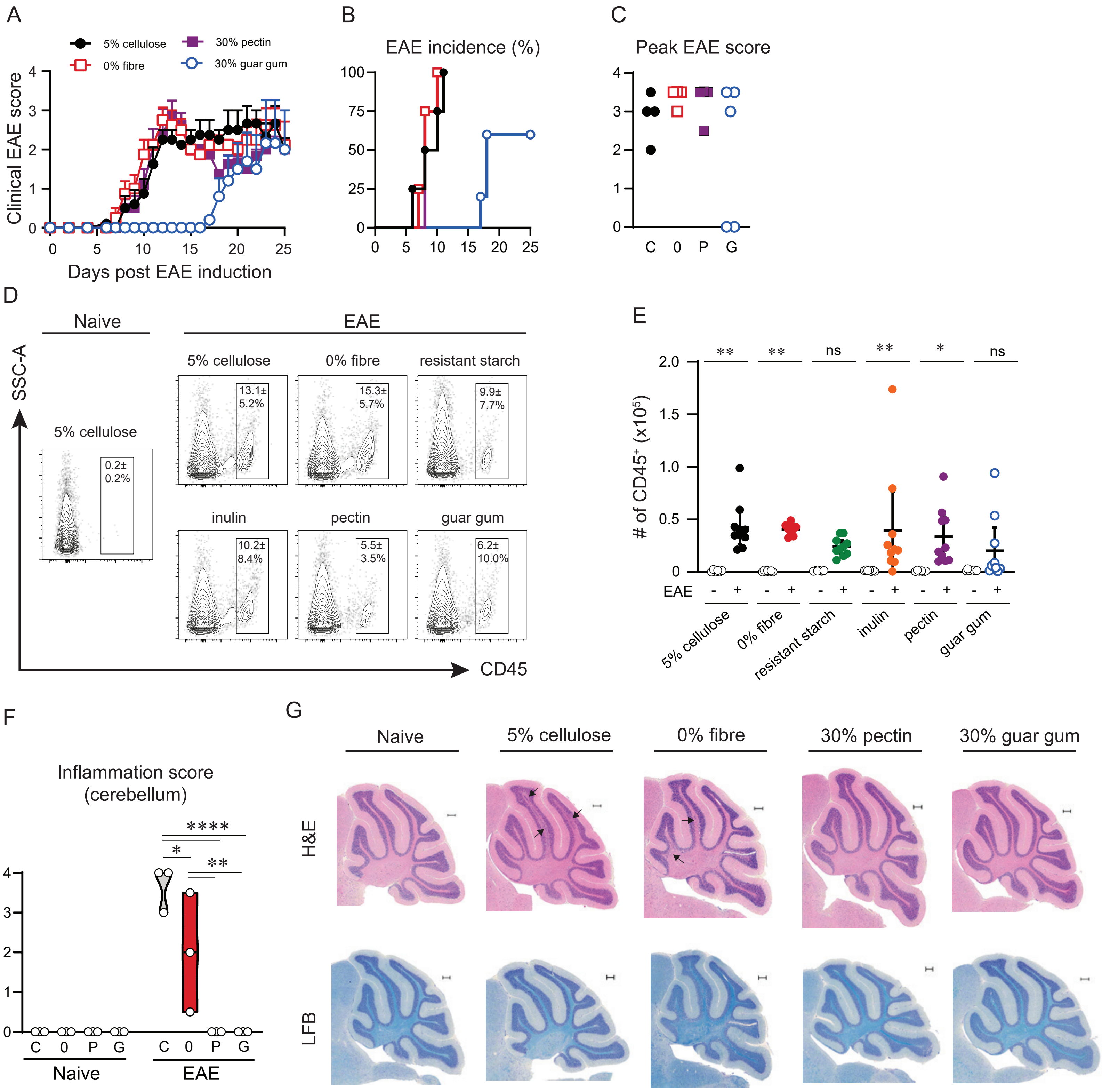

**Supplemental Figure 1. Guar gum supplementation provides long-term protection and limits brain infiltration.** Mice were fed altered-fiber diets for 2 weeks prior to $MOG_{35-55}$ immunization. **A)** Clinical EAE scores of mice on each diet (n=4-5 per group) until d25 post-EAE. Data shown as mean ± SEM. **B)** Percent EAE incidence per day post-EAE induction. **C)** Peak EAE score reached for each individual mouse over the course of EAE. **D)** Concatenated flow plots from one representative experiment of brain-infiltrating $CD45^{hi}$ cells. Data shown mean ± SD for shown experiment of proportion of live cells in the brain. **E)** Quantification of infiltrating $CD45^{hi}$ cells in the brain of naïve and EAE-induced mice for each diet. Data shown as mean ± 95% confidence interval. Statistics by multiple Mann-Whitney tests with Holm-Šídák multiple comparisons test. **F)** Quantification of cellular infiltration in the cerebellum as an inflammation score. Statistics by 2-way ANOVA with Tukey's multiple comparisons test. **G)** Representative images of cerebellum histology of EAE mice by hematoxylin and eosin (H&E) and serial section stained with luxol fast blue (LFB). Naïve sample from 5% cellulose-fed mouse without EAE. Arrows depict regions of cellular infiltration. Scale bar = 200 μm.

### Supplemental Figure 2

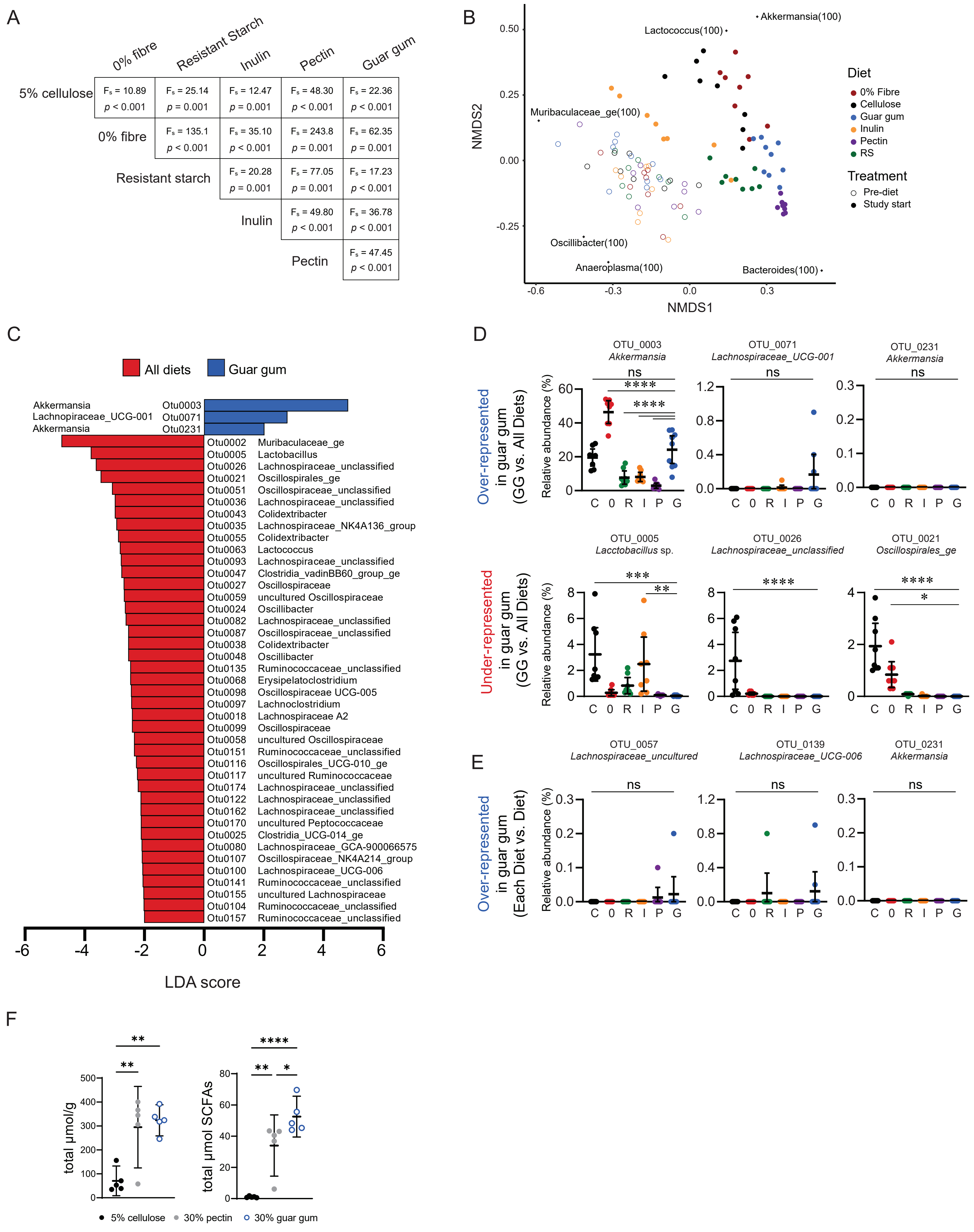

**Supplemental Figure 2. Guar gum does not uniquely change the microbiota compared to other high-fiber diets.** Mice were fed altered-fiber diets for 2 weeks. **A)** AMOVA table of PCoA in Fig 2A. **B)** NMDS of Bray-Curtis dissimilarity based on OTU relative abundance. Highly significant correlates ($R^2$>0.5) between ordination and OTU abundances identified with EnvFit function shown as representative points. **C)** LEfSe analysis of guar gum vs. all diets, depicting OTUs over-represented (blue) in guar gum, and those over-represented in the All Diets group (red). LDA>2, p<0.05. **D)** Visualization of select OTUs identified as over- and under-represented in guar gum by LEfSe (guar gum vs. All Diets). **E)** Visualization of select OTUs identified as over-represented in guar gum by LEfSe (each diet vs. each diet). **F)** Total short chain fatty acids in the cecum per gram of cecal weight (left) and total within each cecum (right).

Data represented as mean ± 95% confidence interval. Statistics by one-way ANOVA with Tukey's multiple comparisons test (B, D-F).

### Supplemental Figure 3

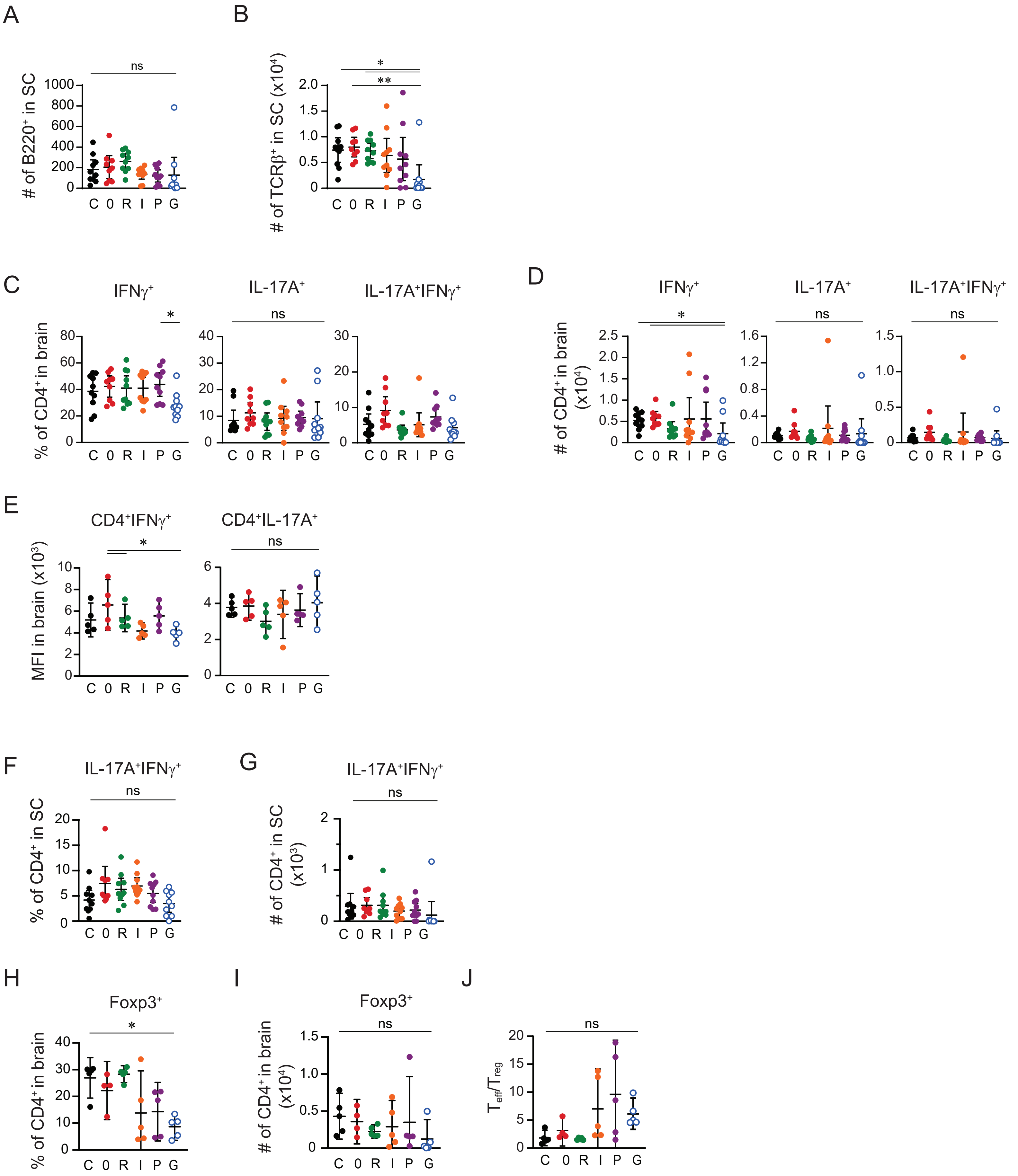

**Supplemental Figure 3. Reduced infiltration of T cells and Th1s into the CNS following guar gum supplementation.** Mice were fed altered-fiber diets for 2 weeks prior to EAE induction. **A)** Number of B220$^+$ B cells in the spinal cord (SC). **B)** Number of TCR$\beta^+$ T cells in the SC. **C)** Proportion of IFN$\gamma^+$, IL-17A$^+$, and IL-17A$^+$IFN$\gamma^+$ CD4$^+$ T cells in the brain. **D)** Proportion of IFN$\gamma^+$, IL-17A$^+$, and IL-17A$^+$IFN$\gamma^+$ CD4$^+$ T cells in the brain. **E)** Median fluorescent intensity (MFI) of IFN$\gamma$ and IL-17A in IFN$\gamma^+$CD4$^+$ T cells and IL-17A$^+$CD4$^+$ T cells in the brain. **F)** Proportion of IL-17A$^+$IFN$\gamma^+$ CD4$^+$ T cells in the spinal cord. **G)** Number of IL-17A$^+$IFN$\gamma^+$ CD4$^+$ T cells in the spinal cord. **H)** Proportion of Foxp3$^+$ CD4$^+$ T cells in the brain. **I)** Number of Foxp3$^+$CD4$^+$ T cells in the brain. **J)** Ratio of cytokine-expressing (IFN$\gamma^+$, IL-17A$^+$, IFN$\gamma^+$IL-17A$^+$) CD4$^+$ T cells to Foxp3-expressing CD4$^+$ T cells in brain.

Data represented as mean ± 95% confidence interval. Statistics by one-way ANOVA with Tukey's multiple comparisons test (A-C,E,F,H,J) or Kruskal-Wallis with Dunn's multiple comparisons test (D,G,I).

### Supplemental Figure 4

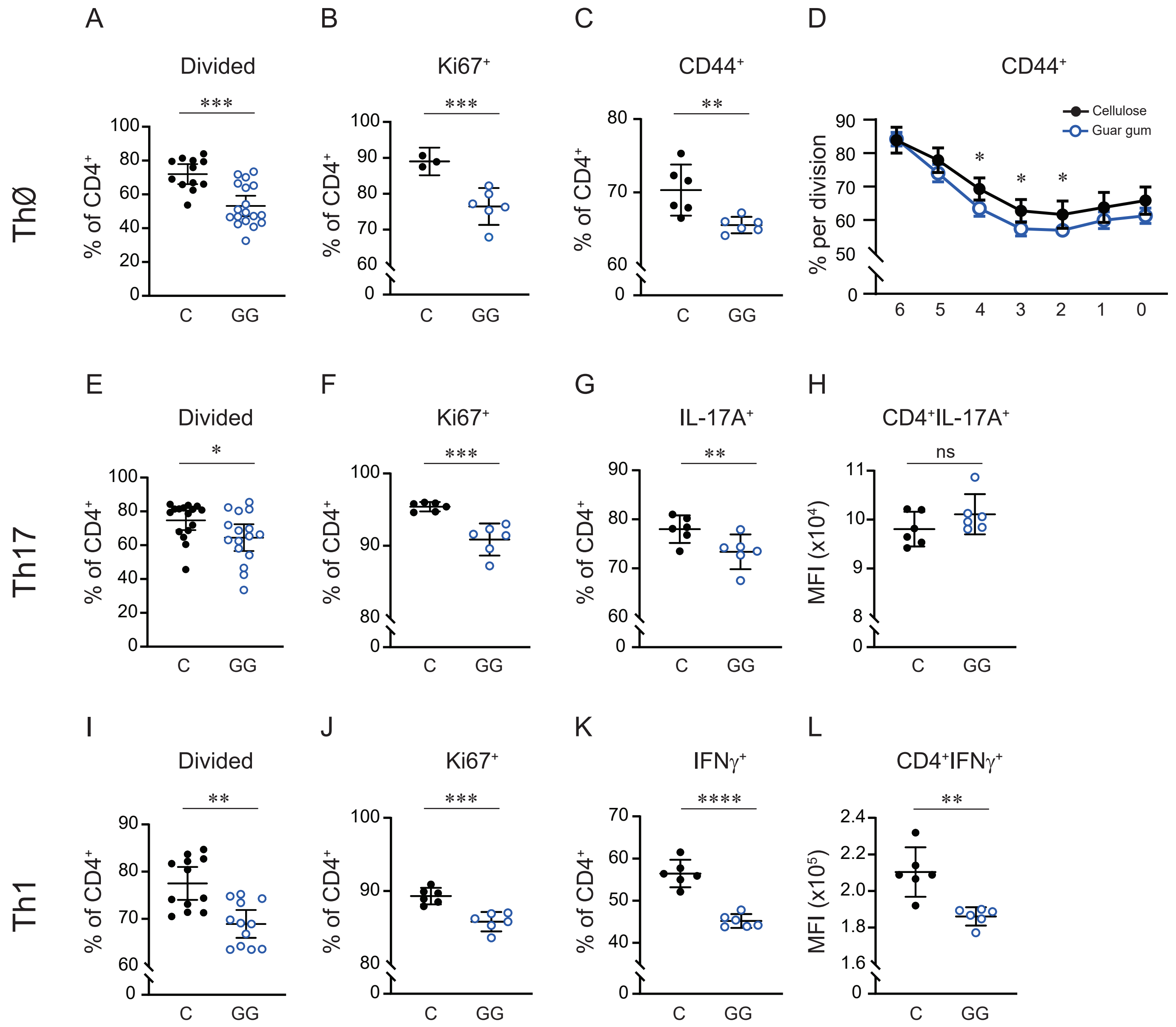

**Supplemental Figure 4. Reduced activation and polarization towards Th1 in guar gum cells is T cell-intrinsic.** $CD4^+$ T cells isolated from spleens of naïve cellulose (C)- or guar gum (GG)-fed mice, stained with CFSE, and cultured with $\alpha$-CD3/$\alpha$-CD28-coated beads for 96 hours in blank culture media (ThØ) **(A-D)**, in Th17-skewing conditions **(E-H),** or in Th1-skewing conditions **(I-L). A,E,I)** Proportion of $CD4^+$ T cells that divided at least once. **B,F,J)** Proportion of $CD4^+$ T cells expressing Ki67. **C)** Proportion of $CD4^+$ T cells in ThØ conditions expressing CD44. **D)** Proportion of $CD4^+$ T cells expressing CD44 at each stage of cell division in ThØ conditions. **G)** Proportion of $CD4^+$ T cells in Th17 conditions expressing IL-17A by flow cytometry. **H)** Median fluorescent intensity (MFI) of IL-17A in $CD4^+$IL-$17A^+$ T cells in Th17 conditions. **K)** Proportion of $CD4^+$ T cells in Th1 conditions expressing IFN$\gamma$ by flow cytometry. **L)** MFI of IFN$\gamma$ in $CD4^+$IFN$\gamma^+$ T cells in Th1 conditions.

All data shown as mean ± 95% confidence interval. Stats by Mann-Whitney test (A,E,I), unpaired t-test (B-C, F-H, J-L) or 2-way ANOVA with Šídák's multiple comparisons test (D).

### Supplemental Figure 5

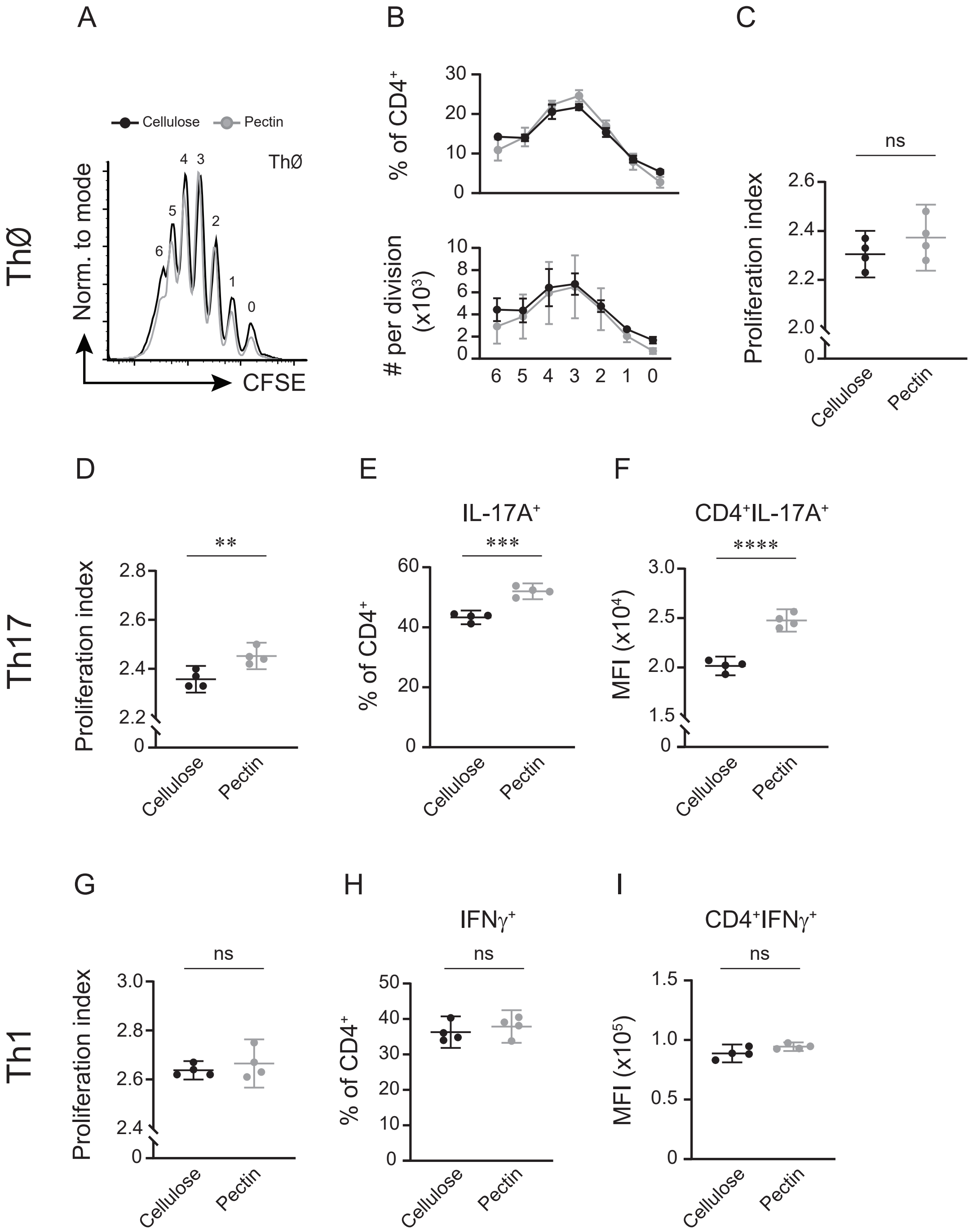

**Supplemental Figure 5. Pectin-derived CD4$^+$ T cells display similar activation and polarization as cellulose-derived cells.** CD4$^+$ T cells isolated from spleens of naïve cellulose- or pectin-fed mice, stained with CFSE, and cultured with $\alpha$-CD3/$\alpha$-CD28-coated beads for 96 hours in blank culture media (ThØ) **(A-C)**, in Th17-skewing conditions **(D-F),** or in Th1-skewing conditions **(G-I). A)** Representative histograms of CFSE dilution in ThØ conditions. **B)** Quantification of proliferation by the proportion (top) and number (bottom) of CD4$^+$ T cells in each division. **C, D, G)** Proliferation index (total number of divisions / cells that went into division) of CD4$^+$ T cells. **E)** Proportion of CD4$^+$ T cells in Th17 conditions expressing IL-17A by flow cytometry. **F)** Median fluorescent intensity (MFI) of IL-17A in CD4$^+$IL-17A$^+$ T cells in Th17 conditions. **H)** Proportion of CD4$^+$ T cells in Th1 conditions expressing IFN$\gamma$. **I)** MFI of IFN$\gamma$ in CD4$^+$IFN$\gamma^+$ T cells in Th1 conditions. Data shown as mean ± 95% confidence interval. Stats by unpaired t-tests.
